# Magnetosome organelles are organized through interactions between McaA and McaB that alter the dynamics of the bacterial actin-like protein MamK

**DOI:** 10.1101/2025.11.10.687720

**Authors:** Yein Ra, Yuanyuan Pan, Tansy Chen, Kayla Dixon, Azuma Taoka, Arash Komeili

## Abstract

Magnetotactic bacteria (MTB) are a group of Gram-negative species that produce a lipid-bounded organelle, the magnetosome, in which a magnetic crystal is biomineralized. MTB use magnetosomes to align with the geomagnetic field for improved navigation of their environment. To optimize this alignment, these species organize their magnetosomes in a linear fashion using a handful of factors including the bacterial actin-like protein MamK. Despite these shared features, there is a broad diversity of species-specific linear magnetosome chain arrangements within MTB, but the molecular mechanisms behind these phenotypic variations are unclear. Recently, genetic analyses showed that two proteins McaA and McaB interface with the chain organization machinery of *Magnetospirillum magneticum* AMB-to arrange magnetite crystals in a series of subchains rather than the single cohesive chains found in closely related MTB. Here, we use *in vivo* co-immunoprecipitation in AMB-1 to demonstrate protein-protein interactions between McaA, McaB, and MamK. Experiments with McaA truncation mutants and conditional control of McaB localization determined that McaA-McaB interactions are dependent on amino acids 530-665 of McaA and McaB localization to the magnetosome chain. We further show that disrupting the McaA-McaB interaction alters the spatial dynamics of MamK *in vivo* We present a model in which protein-protein interactions between McaA, McaB, and MamK drive changes in MamK behavior to establish AMB-1’s magnetosome chain organization.

**IMPORTANCE:** Magnetosomes model for understanding the cell biology of bacterial organelles and the mechanisms of bacterial cell organization. MamK, one of the best-studied bacterial actins, is a notable player in magnetosome chain assembly. This work explores how MamK dynamics are altered by two potential bacterial actin binding proteins, McaA and McaB. This system illustrates how changes in bacterial cytoskeleton regulation result in different organization of subcellular compartments. The conclusions from this research also have implications for understanding the broader evolutionary strategies for regulation of actin-like proteins and compartment organization diversification in bacteria.

## INTRODUCTION

Similar to eukaryotes, many bacterial species produce one or more organelles that carry out a range of biochemical and behavioral activities (1–4). One of the best studied bacterial organelles is the magnetosome (5). Produced by magnetotactic bacteria (MTB), these organelles start as a lipid-bounded compartment wherein a magnetic crystal of magnetite (Fe_3_O_4_) or greigite (Fe_3_S_4_) is biomineralized (6, 7). Magnetosomes allow MTB to align with the Earth’s magnetic field, which is advantageous for navigating their aquatic environment with greater efficiency. Because MTB are typically motivated to find favorable oxygen environments, this behavior is termed magnetoaerotaxis (5). The ability to form magnetosomes makes MTB attractive bacterial system to study biomineralization, organelle biogenesis, and compartment localization.

Magnetosomes exemplify how the function of an organelle can be closely tied to its subcellular localization. A prerequisite to magnetoaerotaxis is the organization of magnetosomes into a chain; various MTB mutants that produce magnetosomes but fail to organize them into a linear chain have dramatically lowered abilities to align with and navigate along a magnetic field (8–12). In the model MTB strains *Magnetospirillum magneticum* AMB-1 (hereafter AMB-1) and *Magnetospirillum gryphiswaldense* MSR-1 (hereafter MSR-1), the magnetosome chain has two important characteristics: it follows the positive curvature of the cell and is continuous, features that are controlled by MamY and MamK, respectively (13, 14). AMB-1 and MSR-1 maintain a linear magnetosome chain in a spiral-shaped body by positioning its chain along the positive cell curvature, thereby ensuring that the chain is parallel to the axis of cell motility. MamY assembles into a scaffold at the positive cell curvature onto which magnetosomes attach through a second protein, MamJ (14). The magnetosome chain is made continuous (i.e. there are no large gaps between magnetosomes) by MamK. MamK is a bacterial actin and in its absence the magnetosome chain is fragmented, not centered, and unevenly segregated during cell division (13, 15–19). Like eukaryotic actin, monomers of MamK polymerize into filaments when bound to ATP (20, 21). Such MamK filaments are observed in AMB-1 and MSR-1 using cryo-electron tomography (9, 13). While within the filament, MamK monomers hydrolyze their ATP into ADP, favoring dissociation from the filament (20). The ATP-dependent polymerization and depolymerization of monomers are required for the dynamic quality and function of MamK. When MamK dynamics are perturbed by disrupting its ATPase activity, both chain assembly and magnetosome segregation are severely hindered (17). MamJ, and its paralog LimJ, connect magnetosomes to MamK filaments and are required for MamK dynamics in AMB-1 (9, 16, 22). MamJ is among the few known accessory factors of a bacterial actin, alongside ParR, AlfB, Alp7R, and others which partner with distinct actin-like proteins to carry out diverse functions (23–25). Identifying additional proteins that regulate MamK can provide further insights to how bacterial cytoskeleton can be controlled.

Recently, two additional proteins that alter MamK behavior have been identified. In AMB-1 strains missing *mcaA* and *mcaB* MamK dynamics changed when observed by fluorescence recovery after photobleaching (FRAP). Additionally, McaA and McaB are required for AMB-1’s magnetosome chain organization. At first glance with transmission electron microscopy (TEM), the WT AMB- magnetosome chain appears fragmented (Fig. 1A). Imaging with cryo-electron tomography, however, reveals that the gaps between crystal subchains are populated with magnetosome membranes that have not yet initiated crystal biomineralization (10). Thus, AMB-magnetosomes organized in subgroups of crystal-containing magnetosomes or empty magnetosomes that together form a continuous chain. In the absence of *mcaA mcaB* or both, the mutant chain resembles that of WT MSR-1, where magnetite crystals are continuous and at midcell while empty magnetosomes are at the chain periphery (Fig. 1A, B) (9, 10, 14). This has collectively led to the hypothesis that McaA and McaB create magnetosome chain with crystal subchains by influencing MamK behavior (10). Investigating how the Mca proteins contribute to magnetosome chain organization will resolve the mechanism behind how AMB- and MSR-1, two organisms that share ∼96% identity in their 16S rRNA genes (26), pattern their magnetosomes in a strikingly different manner. Currently, how McaA and McaB work together at the protein level, and whether they directly affect MamK, is unknown.

**FIG 1:**
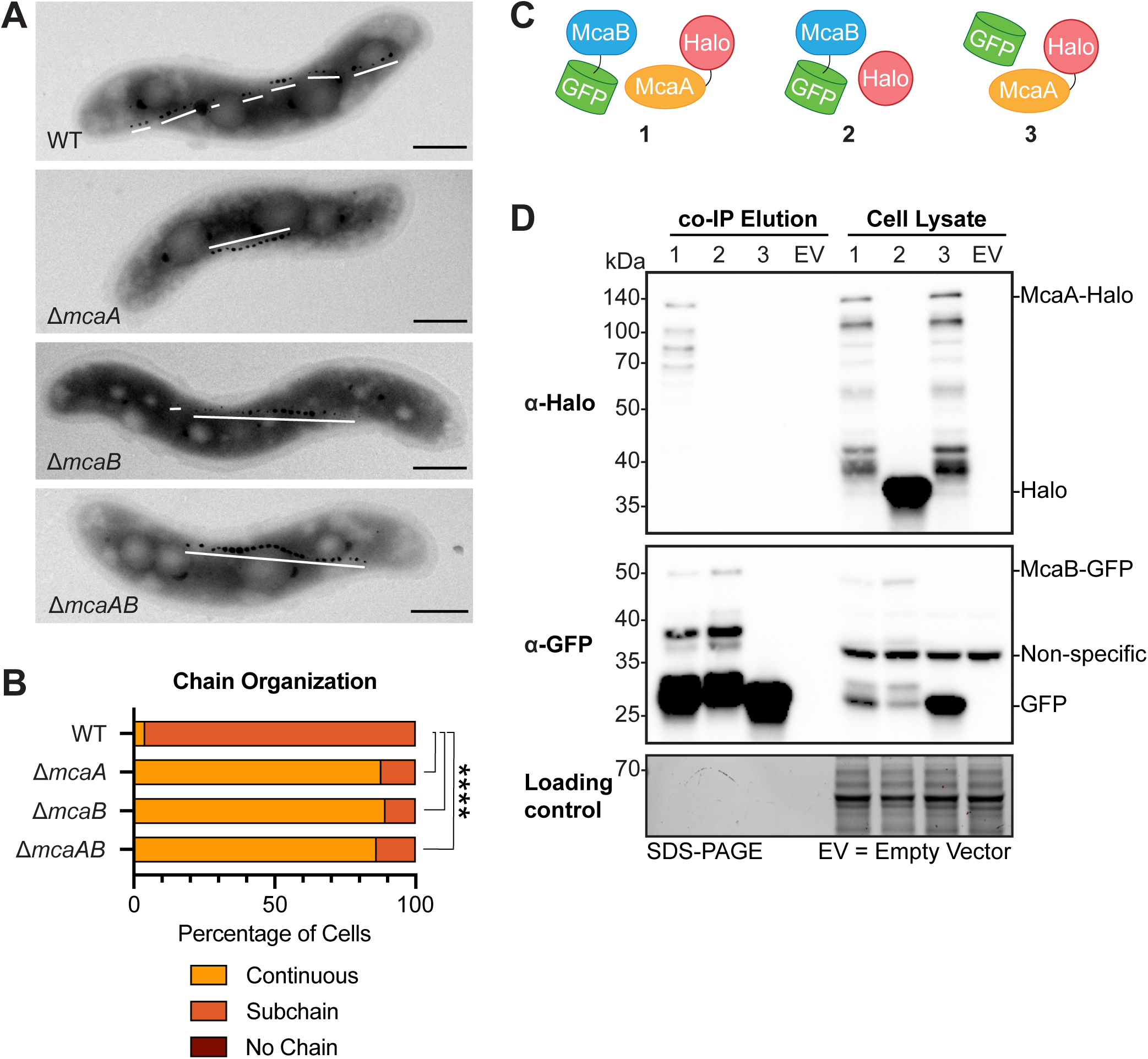
McaA and McaB contribute to magnetosome organization and interact in vivo. (**A**) Transmission electron microscopy images of a representative cell for AMB-1 WT, Δ*mcaA*, Δ*mcaB*, and Δ*mcaAB* showing the magnetosome chain. White lines are drawn to show the placement of crystal-containing magnetosomes. The WT magnetosome chain has many subchains of crystal-containing magnetosomes while Δ*mcaA*, Δ*mcaB*, and Δ*mcaAB* do not. Scale bar = 0.5μm. (**B**) Quantification of magnetosome chain organization using TEM images. Cells were categorized based on their magnetosome chain phenotype and the x-axis represents the percentage of cells that displayed the indicated chain organization. WT n = 109, Δ*mcaA* n = 168, Δ*mcaB* n = 165, Δ*mcaAB* n = 75. Fisher’s exact test was used to determine p-values (**** P < 0.0001). (**C**) Schematics of the combination of proteins used for the *in vivo* co-IP in (D). (**D**) Western blots of elutions from co-IP with GFP-Trap agarose beads and cell lysates illustrating that McaA-Halo and McaB-GFP interact, and this interaction is not driven by their tags. The co-IP was done using the Δ *mcaAB* strain. The loading control is a stain-free SDS-PAGE gel of the same samples.

Here, we discovered that McaA, McaB, and MamK form protein-protein interactions with each other. The McaA-McaB interaction is mediated through a conserved region on McaA, and primarily occurs when McaB is localized to the magnetosome chain. Disrupting the McaA-McaB interactions by truncating McaA or altering McaB localization is accompanied by a shift n MamK dynamics. Altogether, the protein-protein interactions of McaA, McaB, and MamK affect MamK dynamics and ultimately organize magnetic crystals into subchains in AMB-1.

## RESULTS

### McaA and McaB interact with each other

The genetic studies of *mcaA* and *mcaB* suggest that there is a machinery dedicated to producing the crystal subchain magnetosome organization (10). To gain insight on the mechanism behind this magnetosome positioning system, we set out to determine which proteins interact with McaA and McaB *in vivo* in AMB-1. We separately expressed GFP-tagged McaA and McaB on replicative plasmids in AMB-1, which have previously been shown to complement their respective gene deletions (10), and used GFP-trap agarose beads for co-immunoprecipitation paired with mass spectrometry (IP-MS). The experiment was also performed with GFP alone to determine which interactions were due to the tag or non-specific binding to the agarose beads (Supplementary Materials File 2).

We first attempted to identify other magnetosome proteins involved in magnetosome chain organization by looking for proteins that co-eluted with both McaA-GFP and McaB-GFP. The McaB-GFP IP-MS results were poor due to low protein abundance, but we still identified peptides from four magnetosome proteins: MamA, MamQ, MamJ, and Mms6 (Supplementary Materials File 2). MamA and MamQ were also identified in the IP-MS with McaA-GFP. To assess whether MamA has a role in magnetosome chain organization, *mamA* deletion strain was imaged with TEM No deviation in magnetosome chain organization was observed (Fig. S1A, B), suggesting that MamA is not necessary for the crystal subchain phenotype. Because AMB-Δ*mamQ* does not produce magnetosomes, magnetosome arrangement could not be assessed in this strain.

The IP-MS results from McaA-GFP also revealed that McaA and McaB may interact in AMB-(Supplementary Materials File 2). To verify this result, the co-immunoprecipitation (co-IP) using GFP-trap beads was repeated with a Δ*mcaAB* strain expressing McaB-GFP and McaA-Halo. These tagged proteins can complement their respective deletion when assessed by TEM and coefficient of magnetism (C_mag_) measurements (Fig. S2), which measures the degree of magnetic field alignment by MTB cells in a liquid culture (7). The elution was probed for the presence of both proteins using Western blotting. Both proteins co-eluted (Fig. 1C, D; Fig. S3A), indicating that indeed, McaA and McaB interactions in AMB-1. Co-IPs with McaB-GFP/Halo and GFP/McaA-Halo pairs expressed in AMB-Δ*mcaAB* confirmed that the GFP and Halo tags, or nonspecific interactions with beads, were not responsible for the interaction (Fig. 1D, Fig. S3B). Samples containing McaA-Halo almost always had a series of smaller bands in the anti-Halo Western blot (Fig. 1D). These appear to be specific to McaA-Halo, since they are not present in samples expressing other Halo-tagged proteins (Fig. 1D, Fig. S3B, Fig. S13C). These bands could represent McaA-Halo degradation products be result of post-translational processing of McaA-Halo. In the anti-GFP blots of McaB-GFP samples, there is a strong signal from a protein that is around the size of GFP (26.8kDa). In the *mcaB-gfp* fusion, the start codon of GFP was not removed. Thus, there might be an internal ribosome binding site that drives the translation of free GFP. Given that our negative control co-IP with free GFP and McaA-Halo does not detect interaction (Fig. 1D, Fig. S3B), it is unlikely that this free GFP is driving the interaction in the McaA-Halo/McaB-GFP co-IP.

Having observed McaA-McaB interactions in AMB-1, we investigated whether the nteraction between them was direct by using purified proteins. Because McaA and McaB are both predicted to contain transmembrane domains, only their soluble, cytoplasmic portions (McaA_cyto_ aa400-776; McaB_cyto_ aa28-219) were purified. To that end, we constructed McaA_cyto_ with an N-terminal Strep-tag and McaB_cyto_ with an N-terminal maltose binding protein (MBP) tag and purified them using affinity chromatography and size exclusion chromatography (Fig. S4A). We performed *in vitro* pull downs with Strep-McaA_cyto_ and MBP-McaB_cyto_ using streptavidin beads. If there is an interaction between them, then both proteins should be found in the elution fraction. However, the elution only contained Strep-McaA_cyto_ (Fig. S4B). Similarly, bacterial two hybrid assays did not detect an interaction between full-length McaB and full-length McaA McaA_cyto_ (10) (Fig. S7B). It is possible that the interaction between the two proteins requires proper localization to the membrane. Alternatively, the McaA-McaB interaction observed *in vivo* might be bridged by one or more proteins. To test whether any magnetosome proteins fulfills this role, we attempted to repeat the McaB-GFP/McaA-Halo co-IP n AMB-ΔMAI, a strain missing the magnetosome island (MAI) that encodes the majority of AMB-1 magnetosome genes (27). However, McaB-GFP is expressed poorly or unstable in this strain, as indicated by a weak signal in the lysate anti-GFP Western blot (Fig. 2C, Fig. S6A). Thus, we were unable to assess the interaction in this background.

**FIG 2:**
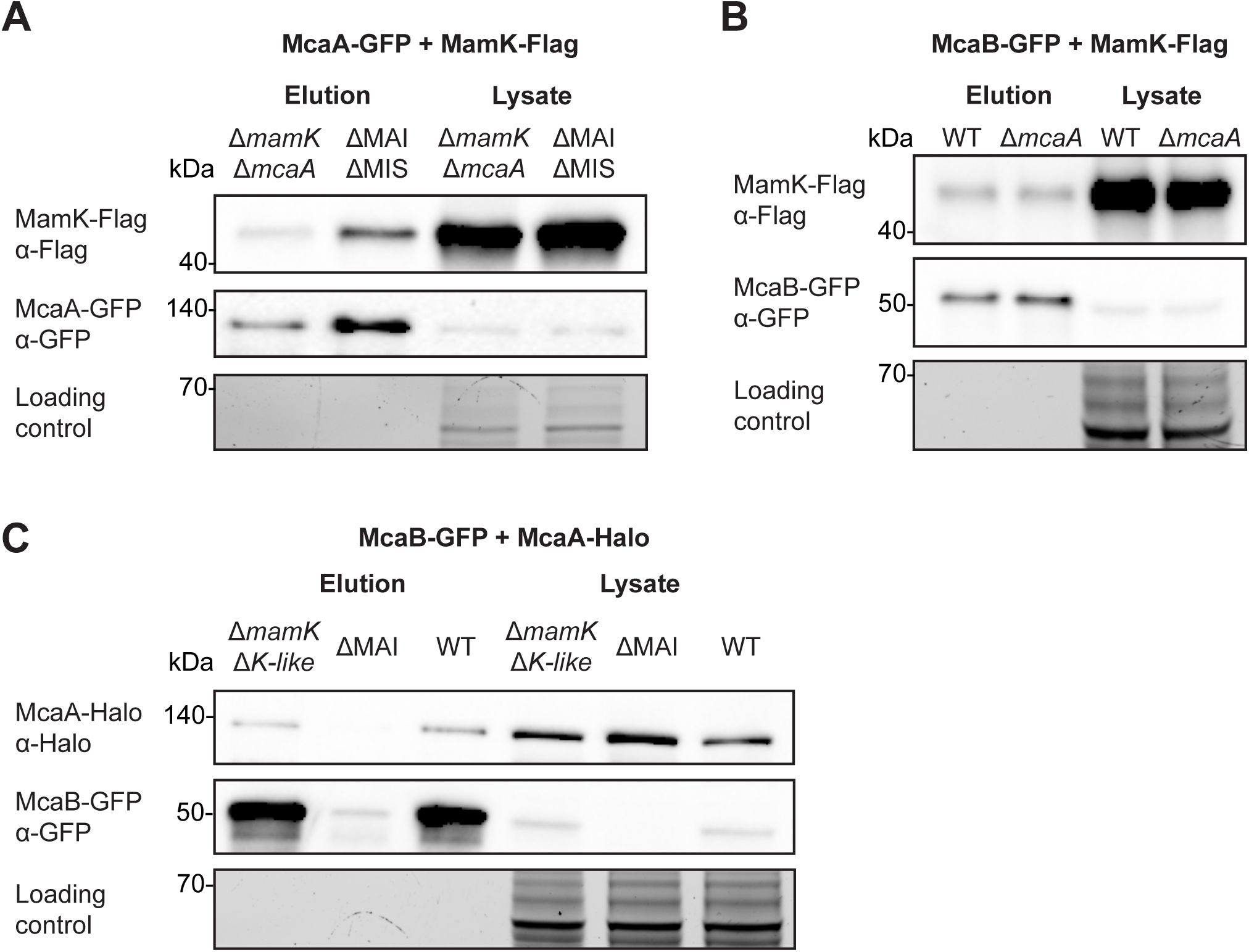
MamK interacts with both McaA and McaB *in vivo*. (**A**) Western blots of a co-IP with McaA-GFP and MamK-Flag done in both Δ*mamK*Δ*mcaA* and ΔMAIΔMIS. (**B**) Western blots of a co-IP with McaB-GFP and MamK-Flag in WT and Δ*mcaA*. (**C**) Western blots of a co-IP with McaB-GFP and McaA-Halo done in Δ*mamK*Δ *mamK-like*, ΔMAI, and WT. In all cases, GFP-Trag agarose beads were used for co-IPs and a stain-free SDS-PAGE serves as the loading control.

### MamK interacts with McaA and McaB *in vivo*

McaA-GFP IP-MS data showed that MamK may also interact with McaA As discussed above, MamK is a bacterial actin that is responsible for assembling magnetosomes into a linear chain (13, 17). Furthermore, MamK dynamics are altered in Δ*mcaA* and Δ*mcaB* strains (10). A co-IP with McaA-GFP and MamK-Flag confirmed the interaction between these two proteins (Fig. 2A, Fig. S5A). Co-IPs with GFP/MamK-Flag and McaA-GFP/Halo-Flag combinations ensured that the observed interactions were not due to the GFP or Flag tags (Fig. S5A). To address whether this interaction is bridged by any other magnetosome protein, including McaB, we assessed whether McaA-GFP and MamK-Flag will interact in ΔMAIΔMIS, a strain missing the magnetotactic islet (MIS) in addition to the MAI (28). We found that MamK-Flag still elutes with McaA-GFP, signifying that the McaA-MamK interaction does not require other magnetosome proteins (Fig. 2A, Fig. S6B).

Since the signal from the McaB-GFP IP-MS experiment was poor, we also considered the possibility that an McaB interaction with MamK might have been missed. We therefore checked for McaB-MamK interactions using co-IP with McaB-GFP and MamK-Flag. Indeed, the two proteins co-eluted (Fig. 2B, Fig. S5B). This interaction is also independent of McaA, as the two proteins still interacted in an *mcaA* deletion background (Fig. 2B, Fig. S6C). The interaction is also not driven by the GFP and Flag tags, since co-IP with GFP/MamK-Flag and McaB-GFP/Halo-Flag in a Δ*mcaB*Δ*mamK* background did not result in co-elution of the protein pairs (Fig. S5B). Finally, we asked whether MamK is necessary for the McaA-McaB interaction. We repeated the McaB-GFP and McaA-Halo co-IP in AMB-Δ*mamK*Δ*mamK-like* double mutant missing both *mamK* and its homolog *mamK-like* that is located within the MIS (28). McaB-GFP was stable in this background and interacted with McaA-Halo (Fig. 2C, Fig. S6A). In summary, we used a series of co-IP experiments to show that McaA, McaB, and MamK all have independent pairwise interactions with one another in AMB-1.

### MamK directly interacts with McaA and McaB

The protein-protein interactions between McaA, McaB, and MamK paired with the previously observed effect of McaA and McaB on MamK dynamics raised the possibility that the Mca proteins are direct regulators of MamK behavior. To investigate direct interactions between these proteins, untagged recombinant MamK was expressed in *Escherichia coli* BL21 and purified using ammonium sulfate precipitation and size exclusion chromatography (Fig. S4A). Purified MamK tested for interactions against recombinant McaA_cyto_ and McaB_cyto_ *in vitro* MamK forms filaments when bound to ATP, which can be separated from MamK monomers using a pelleting assay (Fig. 3B). We used this assay to assess whether McaA or McaB co-pellets with MamK filaments. MBP-McaB_cyto_ co-pelleted with MamK in an ATP-dependent manner while MBP tag alone predominantly remained in the supernatant (Fig. 3B), signifying that McaB_cyto_ directly binds MamK filaments. Strep-McaA_cyto_ pellets on its own with ATP (Fig. 3A). Therefore, we could not use this assay to determine its direct interaction with MamK. We instead turned to bacterial two hybrid assays, which detected an interaction between McaA_cyto_ and MamK (Fig. S7A). This interaction was missed in a previous bacterial two hybrid assay where LB medium was used instead of M63 (10). Thus, MamK interacts with both McaA and McaB directly as observed by pelleting and bacterial two hybrid assays.

**FIG 3:**
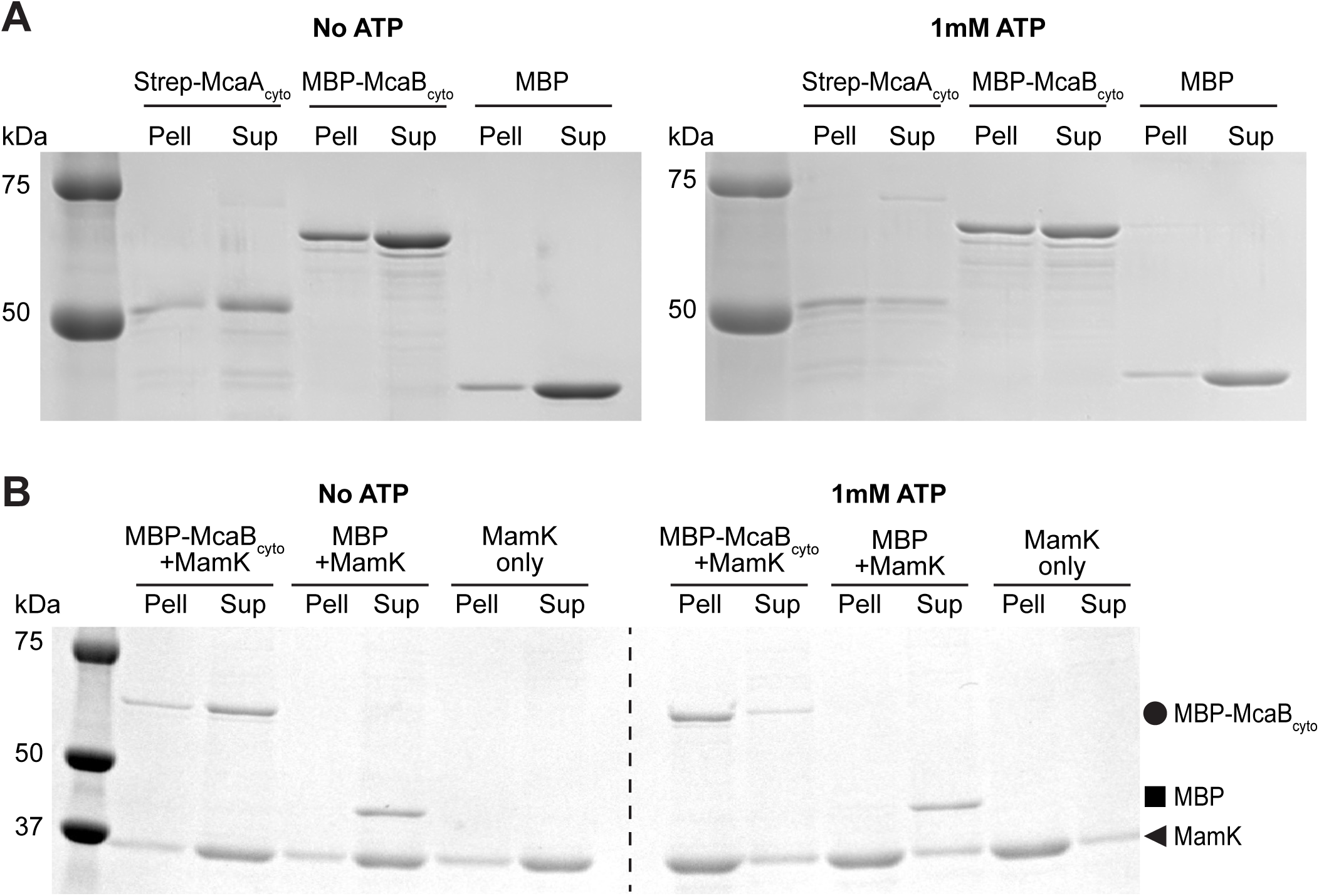
Purified McaB_cyto_ interacts with MamK filaments. (**A**) SDS-PAGE of pelleting assay samples. Proteins were incubated with or without 1mM ATP, then separated into the pellet (Pell) and supernatant (Sup) fractions. (**B**) SDS-PAGE of co-pelleting assay samples. MamK was mixed with either MBP-McaB_cyto_ or MBP and incubated with or without 1mM ATP. The sample was separated into the pellet (Pell) and supernatant (Sup) fractions. MamK polymerizes into filaments in the presence of ATP, demonstrated by the shift into the pellet fraction when incubated with ATP. MBP-McaB_cyto_ shifts into the pellet fraction in the presence of MamK filaments, indicating an interaction between MBP-McaB_cyto_ and MamK. MBP alone remains predominantly in the supernatant in the parallel experiment. MBP-McaB_cyto_ (65.0kDa), MBP (43.2kDa), and MamK (37.6kDa) bands are indicated with a circle, square, and triangle respectively.

### Interaction between McaA and McaB is mediated through a conserved domain of McaA

Next, we asked whether disrupting the McaA-McaB interaction will have an impact on their function *in vivo* To do so, we took a closer look at the predicted secondary structures of the Mca proteins to identify potential domains that mediate their interactions with one another. McaB is a membrane protein that localizes to crystal-containing magnetosomes (10). Analysis with DeepCoil2 revealed a high probability that McaB amino acids 81-112 forms a coiled-coil domain (Fig. S8D) (29). Coiled-coil domains are alpha-helixes that oligomerize by burying their hydrophobic residues, commonly referred to as “a” and “d” position amino acids, against each other. We mapped the “a” and “d” position amino acids predicted from DeepCoil2 onto the AlphaFold-predicted McaB monomer, dimer, and trimer (Fig. S8A-C) (30). We found that in a trimer, the “a” and “d” position residues are indeed predicted to form a hydrophobic core (Fig. S8E).

McaA is a membrane protein that localizes to the positive inner curvature of the cell independently from magnetosomes (10). It has a predicted periplasmic von Willebrand factor type A (VWA) domain at its N-terminus that is necessary for its localization to the positive curvature (Fig. S9A, Fig. S10A) (10). On the C-terminal cytoplasmic side of the transmembrane domain, McaA has a stretch of amino acids from 530 to 665 (named the “530 region”) that is more conserved compared to rest of the C-terminal side (Fig. S9B). It has been shown that McaA without the 530 region (McaAΔ530) is unable to complement the *mcaA* deletion; Δ*mcaA* expressing McaAΔ530-GFP on a plasmid still produces a continuous chain of magnetite crystals despite having the same localization as full-length McaA-GFP (10). McaAΔ530-GFP is also the only truncation made on the cytoplasmic side of McaA that cannot complement the Δ*mcaA* deletion (10). Guided by Alphafold3 structure predictions (30), the 530 region was further divided into three smaller segments and each was removed from McaA-GFP (Fig. S10A). Each GFP-tagged truncation had the same localization pattern as full length McaA-GFP (Fig. S10F). However, none of the three truncated McaA-GFP restored the crystal subchain magnetosome organization when expressed in the *mcaA* deletion, and the C_mag_ of these strains was similar to Δ*mcaA* with an empty vector (Fig. S10C-E). Thus, the entire 530 region is necessary for McaA’s function. We hypothesized that the loss-of-function of McaAΔ530 could be due to loss of interaction with McaB or MamK. To test this, expressed McaB-GFP and McaAΔ530-Halo in Δ*mcaAB* and tested for interaction between the proteins using a co-IP assay. Indeed, McaAΔ530-Halo did not elute with McaB-GFP (Fig. 4A). On the other hand, McaAΔ530-GFP retained its interaction with MamK-Flag in Δ*mcaA*Δ*mamK* and ΔMAIΔMIS strains (Fig. S5A, Fig. S6B). Thus, the 530 region is necessary for McaA function and its interaction with McaB, and the loss-of-function of McaAΔ530 is not due to lack of interaction with MamK.

**FIG 4:**
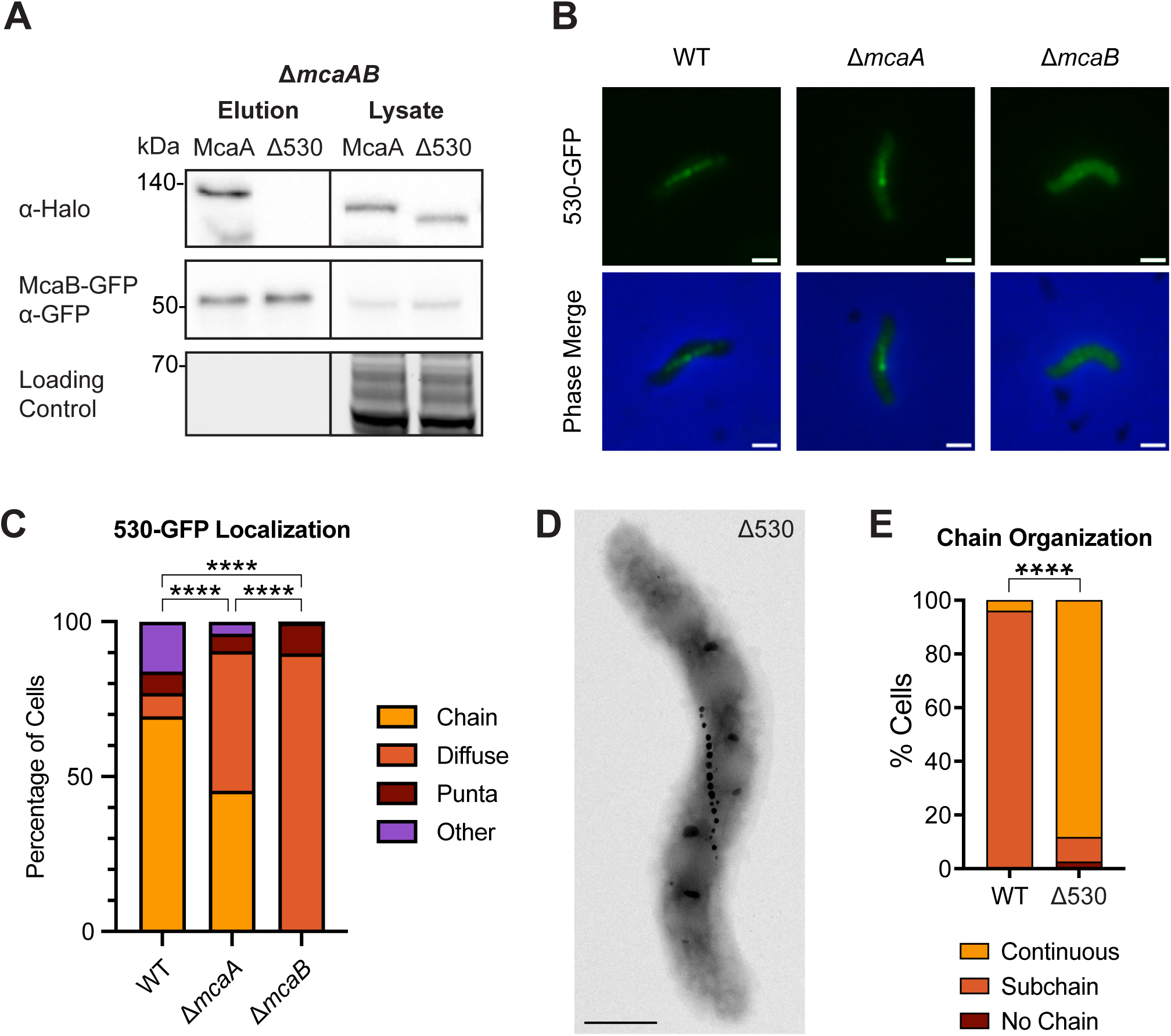
The 530 region of McaA is necessary for interaction with McaB and crystal subchain phenotype. (**A**) Western blots of co-IPs with McaB-GFP and either full-length McaA-Halo or McaA-Halo missing the 530 region (Δ530) in a Δ *mcaAB* strain using GFP-Trap agarose beads. The loading control is a stain-free SDS-PAGE gel. (**B**) The 530 region fragment was tagged with GFP and expressed in WT, Δ*mcaA*, and Δ*mcaB*. Representative cells for each expression strain are shown. Scale bar = 1mm. (**C**) Quantification of 530-GFP localization from fluorescent microscopy images. Cells were categorized based on their localization pattern and the y-axis represents the percentage of cells that displayed the indicated localization. WT n = 390, Δ*mcaA* n = 390, Δ*mcaB* n = 332. Fisher’s exact test was used to determine p-values (**** P < 0.0001). (**D**) Representative TEM image of AMB-1 with the 530 region deleted at the native locus. Scale bar = 0.5μm. (**E**) Quantification of magnetosome chain organization using TEM images. Cells were categorized based on their magnetosome chain phenotype and the y-axis represents the percentage of cells that displayed the indicated chain organization. WT n = 75, Δ530 n = 77. Fisher’s exact test was used to determine p-values (**** P < 0.0001).

To determine whether the 530 region is sufficient for interaction with McaB, we cloned a fragment of McaA spanning this region and tagged it with GFP (530-GFP). We then expressed this construct in WT, Δ*mcaA* and Δ*mcaB* In WT, the majority of cells displayed 530-GFP with a magnetosome chain localization (Fig. 4B, C). This localization is not dependent on full-length McaA; though there increase in cells with diffuse 530-GFP signal, large proportion of the population still had 530-GFP localized to the magnetosome chain (Fig. 4B, C). This magnetosome chain localization is almost completely lost in the absence of *mcaB* the vast majority of Δ*mcaB* cells have a diffuse 530-GFP signal (Fig. 4B, C). One explanation for this is that the 530 region is sufficient to interact with McaB, and this interaction draws 530-GFP to the magnetosome chain.

### McaA 530 region is necessary for WT MamK dynamics

It has been previously shown that MamK dynamics are impacted by McaA and McaB. In prior FRAP experiments with WT AMB-expressing MamK-GFP, the photobleached region was always static and recovered in place (10). In Δ*mcaA* and Δ*mcaB* o the other hand, the photobleached region was motile in a portion of the cells, indicating that MamK dynamics is influenced by the Mca proteins (10). We hypothesized that the interaction between McaA and McaB, which appears critical for magnetosome chain patterning, also contributes to the dynamics of MamK recovery in FRAP.

To test this, we first made an in-frame deletion of the 530 region at the native *mcaA* locus. Like Δ*mcaA* this strain has a magnetosome chain with crystal-containing magnetosomes at the chain middle (Fig. 4D, E), and its C_mag_ is higher than that of WT (Fig. S10B). We then expressed MamK-GFP in WT, Δ*mcaA* and Δ530, and observed MamK dynamics using FRAP. We found MamK-GFP to be dynamic in all strain backgrounds (Fig. 5A, 5C; Fig. S11). We first compared the half-time of fluorescence recovery (t_1/2_), the time it takes for the bleached region to regain 50% fluorescence intensity of the whole filament intensity (Table 1). When examining the entire population of recovered cells across WT, Δ*mcaA* and Δ530, the t_1/2_ did not substantially vary (3.53 ± 3.15 min, 3.02 ± 2.22 min, 3.65 ± 3.67 min, respectively). In most WT cells that recovered, the bleach spot was stationary and recovered in place. However, the bleach spot moved during recovery in about 15% of WT cells observed In a Δ*mcaA* strain, the number of cells with a MamK-GFP moving bleach spot more than doubled (38% of recovered cells observed), consistent with prior observations (10). Similarly, 38% of recovered Δ530 cells exhibited the moving bleach spot pattern (Fig. 5A, B). MamK-GFP behavior thus shifted more towards moving-bleach-spot dynamics in Δ530, as in Δ*mcaA* (Fig. 5C). When we separated the recovered cell population based on MamK recovery behavior (non-moving vs moving bleach spot), we can see that in general, cells with a moving bleach spot recover faster than those with non-moving bleach spots (Table 1). Recovery times were independent of cell length, with the exception of Δ530 cells in which those with non-moving bleach spots showed a slight positive correlation (Fig. S12A). The directions of the moving bleach spots were not consistent from cell to cell; the moving bleach spot moved towards the cell pole in some cells and towards midcell in others (Fig. S12B). Altogether, change in magnetosome organization and altered MamK dynamics correlates with the loss of the McaA-McaB interaction.

**FIG 5:**
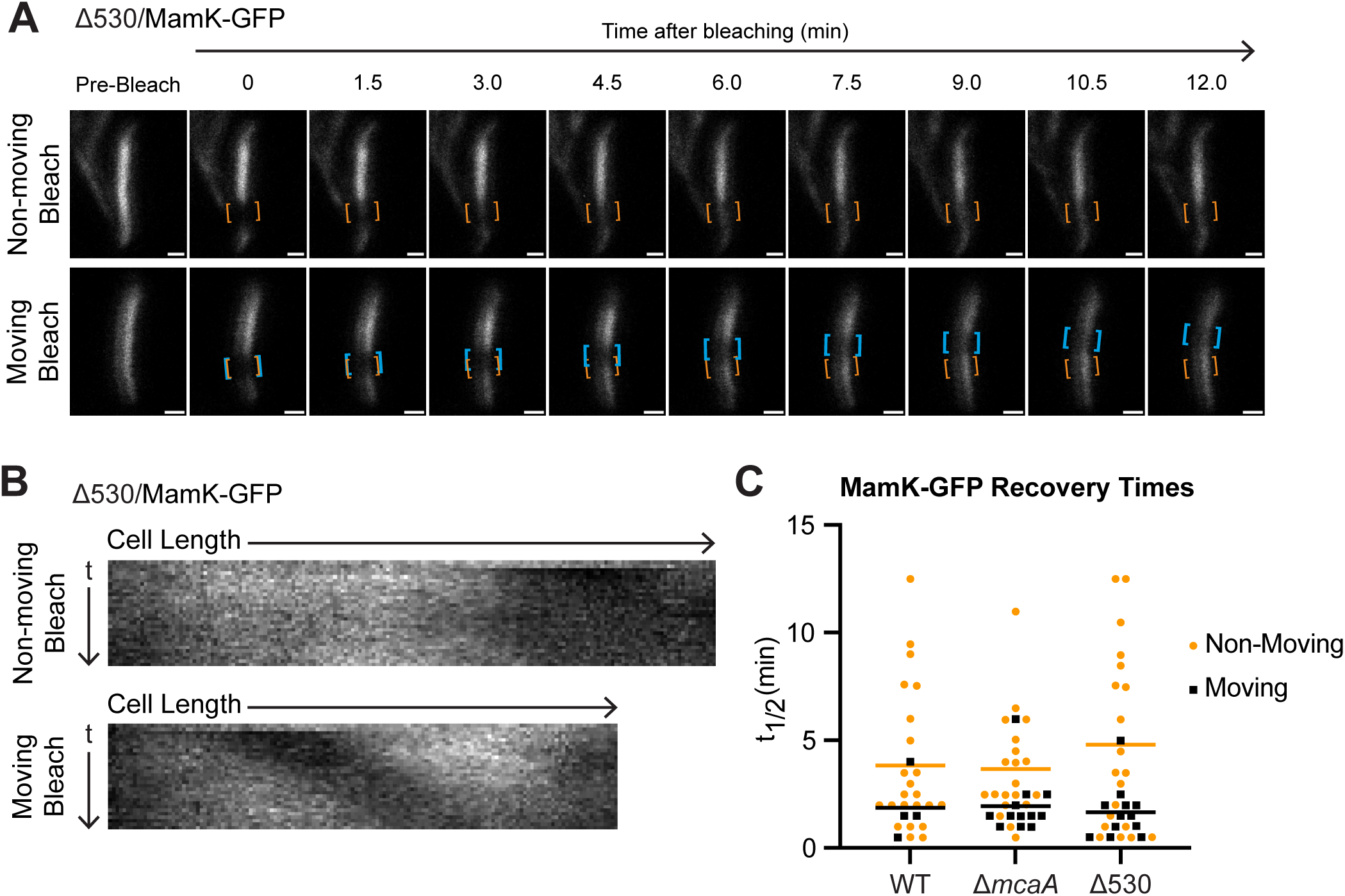
The 530 region is required for WT MamK dynamics. (**A**) Two representative cells from FRAP of MamK-GFP in Δ530 strain. Orange brackets indicate the original bleaching location. Blue brackets track bleach spot movements. The top cell has a bleach spot that does not move while the bottom cell has a moving bleach spot. Scale bar = 0.5 μm. (**B**) Kymographs of the cells in (A). (**C**) MamK-GFP half-time of fluorescence recovery (t_1/2_) across different AMB-1 strains. Each point represents one cell. Yellow circles indicate cells with non-moving bleach spots while black triangles represent those with a moving bleach spot. Cells that did not recover were not included. Bars represent the average recovery half-time of each population.

**Table 1:** MamK-GFP dynamics observed by FRAP.

| Strain | n | % recovered | % moving bleach of recovered | t1/2 average (min) ± SD | | |
| --- | --- | --- | --- | --- | --- | --- |
|  |  |  |  | All cells | Non-moving bleach cells | Moving bleach cells |
| WT/MamK-GFP | 31 | 87 (n = 27) | 15 (n = 4) | 3.53 ± 3.15 | 3.82 ± 3.29 | 1.87 ± 1.50 |
| ΔmcaA/MamK-GFP | 32 | 100 (n = 32) | 38 (n = 12) | 3.02 ± 2.22 | 3.65 ± 2.41 | 1.95 ± 1.37 |
| Δ530/MamK-GFP | 35 | 91 (n = 32) | 38 (n = 12) | 3.65 ± 3.67 | 4.79 ± 4.14 | 1.66 ± 1.24 |
| WT/MamK-GFP, nonbiomin | 32 | 78 (n = 25) | 36 (n = 9) | 5.11 ± 3.56 | 5.06 ± 3.64 | 5.21 ± 3.63 |

### Non-biomineralization conditions affect McaB-GFP localization, McaA-McaB interaction, and MamK dynamics

McaAB-mediated change of MamK dynamics appears to be dependent on McaA-McaB interactions. This led to consider that MamK behavior might be locally controlled based where McaA-McaB interactions occur along the magnetosome chain. We further speculated that McaA-McaB interactions are limited to where the two proteins co-localize To assess the role of localization on McaA-McaB interactions, we perturbed McaB localization to crystal-containing magnetosomes.

McaB’s localization pattern resembles that of another magnetosome protein, Mms6. Mms6 is involved in the biomineralization of the magnetite crystal, and like McaB, is present magnetosomes that contain magnetite (31, 32). Furthermore, Mms6 was identified in the IP-MS with McaB-GFP. Therefore, we hypothesized that McaB is recruited to crystal-containing magnetosomes through an interaction with Mms6. We tested this by comparing McaB-GFP localization in WT and Δ*mms6* At first glance, the McaB-GFP signal seems diffuse in Δ*mms6* compared to WT. However, upon closer inspection, noted that subpopulation of McaB-GFP is still found at the magnetosome chain in most cells (Fig. S13A, B). Additionally, we could not confirm the Mms6-McaB interaction through co-IPs with Mms6-Halo and McaB-GFP in WT AMB-(Fig. S13C). A co-IP of McaB-GFP and McaA-Halo determined that the two proteins still interact in Δ*mms6* (Fig. S13D) and TEM images of AMB-1 Δ*mms6* revealed only a minimal change in magnetosome arrangement in this strain (Fig. S13E, F). Therefore, Mms6 is not necessary for recruitment of McaB to the magnetosome chain, the McaA-McaB interaction, and proper chain formation.

We next turned to previous studies which showed a conditional change in McaB-GFP localization. When AMB-cells are shifted to an iron-limited growth medium, they produce a chain of magnetosome membranes that are devoid of magnetite crystals (7, 27). In previous work, we found that McaB-GFP localization appears diffuse under these non-biomineralizing conditions (10). We first quantified the difference in McaB-GFP localization in both biomineralization and non-biomineralization conditions in WT AMB-1. Indeed, in non-biomineralization growth, the majority of AMB-1 cells show diffuse McaB-GFP signal (Fig. 6A, B). We then tested whether McaB still interacts with McaA when it is no longer localized to the magnetosome chain. Compared to biomineralizing conditions, McaB interactions with full-size McaA are notably reduced in non-biomineralizing conditions (Fig. 6C). Interestingly, shorter McaA-Halo bands still co-eluted with McaB-GFP (Fig. S14A). We speculate that some degradation products of McaA-Halo have lost their transmembrane domain. This soluble McaA-Halo truncation can then interact with the diffuse McaB-GFP. Altogether, McaB localization to the magnetosome chain is a prerequisite for its interaction with McaA.

**FIG 6:**
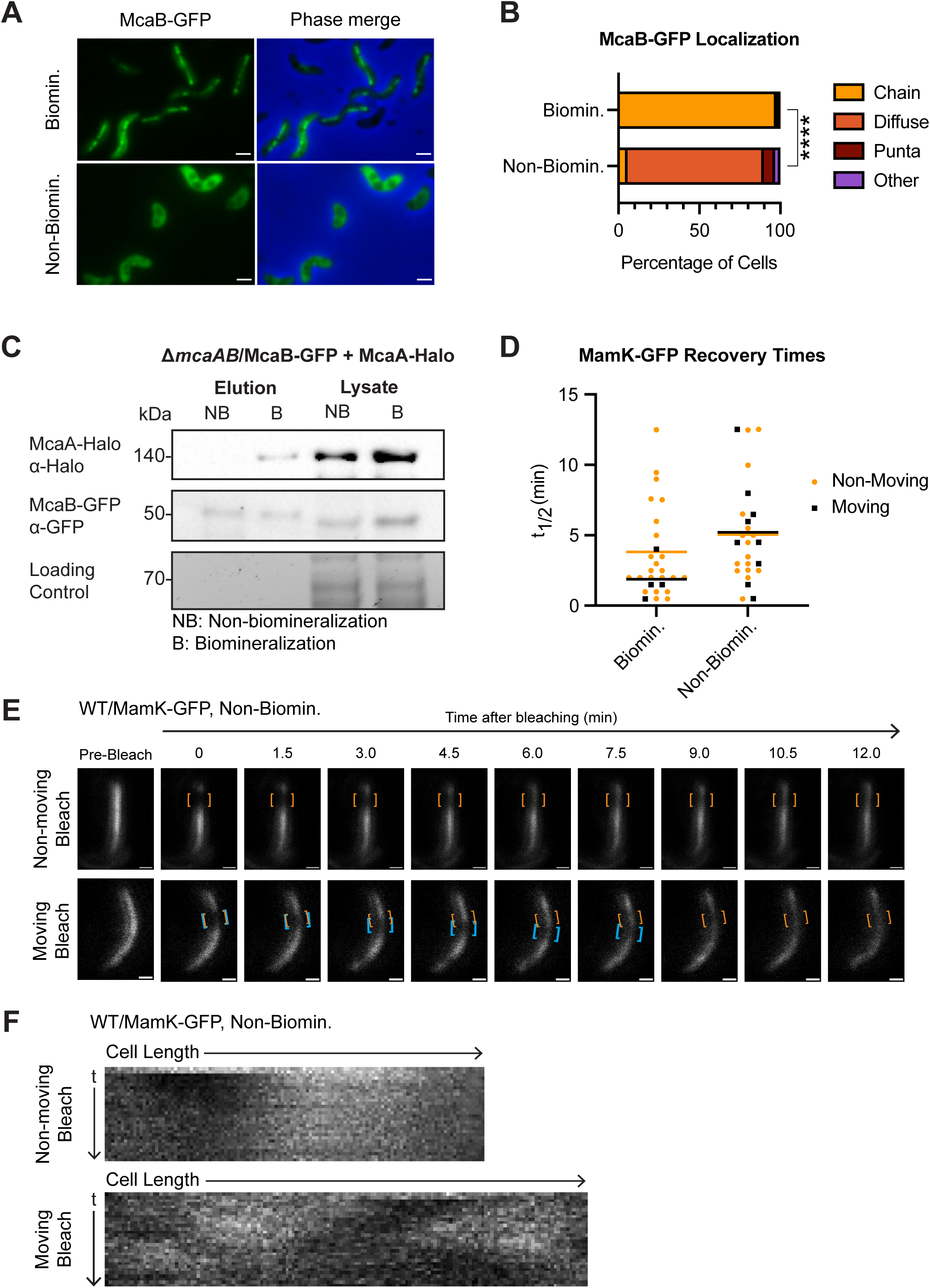
Changes to McaB localization correlates with its ability to interact with McaA and shift in MamK dynamics. (**A**) Representative cells showing McaB-GFP localization in biomineralization and non-biomineralization conditions in the same strain used for co-IP in (C). Scale bar = 1μm. (**B**) Quantification of McaB-GFP localization using fluorescent microscopy images. Cells were categorized based on their localization pattern and the y-axis represents the percentage of cells that displayed the indicated chain organization. Biomineralization n = 845, non-biomineralization n = 4243. Chi-squared test of independence was used to determine p-values (**** P < 0.0001). (**C**) Western blots of co-IP with McaB-GFP and McaA-Halo in non-biomineralization (NB) and biomineralization (B) conditions in a Δ *mcaAB* background using GFP-Trap agarose beads. The loading control is a stain-free SDS-PAGE gel. (**D**) Recovery half-times (t_1/2_) of MamK-GFP in WT AMB-1 under biomineralization (same as WT from Fig. 5C) or non-biomineralization conditions. Each point represents one cell. Yellow dots indicate cells with a non-moving bleach spot while black squares represent those with a moving bleach spot. Cells that did not recover were not included. Bars represent the average recovery half-time of each population. (**E**) Two representative cells from FRAP of MamK-GFP in WT grown under non-biomineralizing conditions. Orange brackets indicate the original bleaching location. Blue brackets track bleach spot movements. The top cell has a bleach spot that does not move while the bottom cell has a moving bleach spot. Scale bar = 0.5μm. (**F**) Kymographs of the same cells depicted in (E).

Since McaA-McaB interactions are missing in non-biomineralizing conditions, we predicted that MamK dynamics would also be impacted. We subjected WT AMB-expressing MamK-GFP to non-biomineralization conditions, then conducted FRAP. We found that in general, MamK-GFP recovery was much slower when cells are grown in non-biomineralization conditions with an average t_1/2_ of 5.11 ± 3.56 min, and 22% of cells did not recover in the observed timeframe (Table 1). The longer recovery times may reflect a broader metabolic constraint from growing in iron-limited medium, demonstrated by the decreased growth observed under non-biomineralization conditions (Fig. S14B). Despite the overall increase in t_1/2_ we observed that 36% of recovering cells displayed the moving-bleach-spot dynamics, similar to Δ*mcaA* and Δ530 (Fig. 6D-F, Table 1). This is consistent with our hypothesis that the effect of McaA and McaB on MamK dynamics is dependent on their ability to interact with each other, which is in part driven by McaB’s localization to the magnetosome chain.

## DISCUSSION

### McaA-McaB interactions drive magnetosome organization through MamK dynamics

In this study, we set out to determine how McaA and McaB produce the subchain organization of magnetosomes in AMB-1. Through both *in vivo* and *in vitro* work, we discovered that McaA and McaB have protein-protein nteractions with each other and the bacterial actin MamK. We further demonstrated that each pairwise interaction occurs independently of the third protein. Disrupting one of these interactions (McaA-McaB) leads to change in magnetosome organization from subchains of magnetite crystals to having singular crystal chain This points to model where network of McaA, McaB, and MamK interactions are required for magnetosome chain organization.

The effect of McaAB on magnetosome organization is closely tied with MamK dynamics. Two types of MamK dynamics were observed through our FRAP imaging: moving-bleach-spot dynamics, where the bleach spot migrated from its original position while recovering, and recovery-in-place dynamics, where the bleach spot recovered without moving. t s challenging to extract the behavior of individual MamK filaments from this data because during FRAP, many overlapping 200nm-sized MamK filaments are simultaneously photobleached (13). Consequently, our interpretation of the FRAP data is focused on bulk MamK filament behavior. We predict that the moving-bleach-spot dynamic is due to a global organization of individual MamK filaments in which the majority of MamK filaments move or polymerize in the direction (Fig. 7A). We currently do not know how the movement is coordinated or how the movement direction is determined. In contrast, the recovery-in-place dynamics is perhaps a result of ndividual MamK filaments moving or polymerizing in either direction along the cell length (Fig. 7A). Though the directionality of individual filaments might be regulated locally, the movement direction is not uniform throughout the entire population of MamK filaments. Interestingly, MamK dynamics are not identical within an entire population. Even in populations that had more cells with moving-bleach dynamics, the majority of cells still showed recovery-in-place dynamics. This implies that there are additional controls to MamK dynamics in AMB-1 that are left to be discovered.

**FIG 7:**
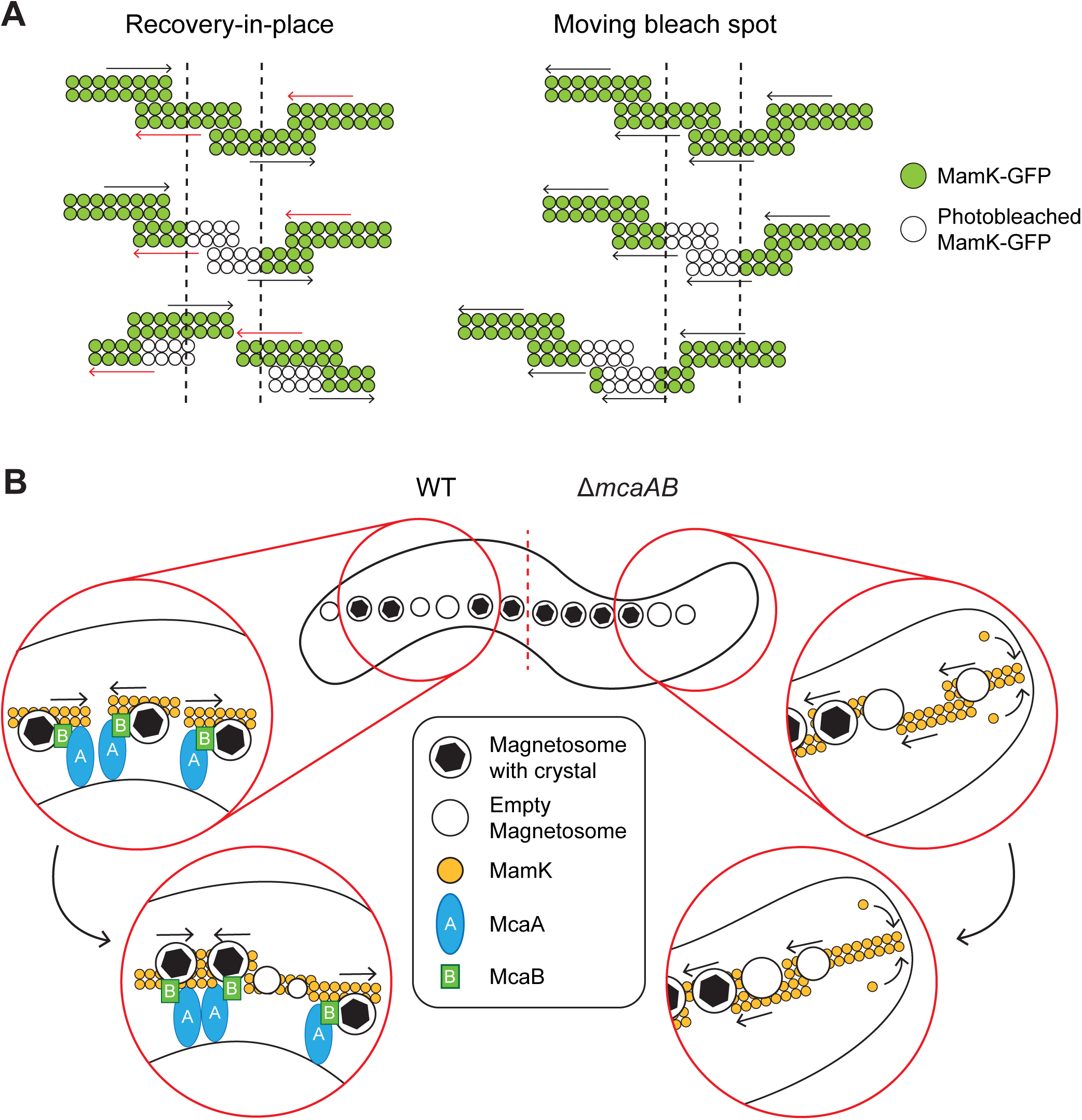
Model for how the magnetosome chain organization is established in WT AMB-1. (**A**) Schematic displaying how the dynamics of MamK filaments produced the two observed FRAP behaviors. Green circles depict fluorescent MamK-GFP while white circles represent photobleached MamK-GFP during the FRAP experiments. The dashed lines indicate the location of photobleaching. In situations where the MamK-GFP bleach spot does not move and is recovering-in-place (left), nearby MamK filaments are moving or polymerizing in either direction, represented by the arrows. The bleached filaments may move away while unbleached filaments move into the region, causing the gradual recovery of the bleached location in place. In cells where the bleach spot is moving (right), nearby MamK filaments have coordinated movement in a single direction. The bleach spot also recovers during its movement, likely because the individual filaments undergo polymerization while moving. (**B**) Model for how McaA and McaB produce crystal subchains by altering MamK dynamics in AMB-1. In WT (left), McaA-McaB interactions occur along the magnetosome chain where crystal-containing magnetosomes are present. This allows the McaAB complex to interact with MamK filaments to alter their directionality, resulting in subgroups of crystal-containing magnetosomes. Concentrating crystal-containing magnetosome into subchains creates space throughout the cell for new magnetosomes to form. In the absence of *mcaAB* (right), MamK filaments move in a single direction, pushing magnetosomes to the midcell. The oldest magnetosomes - most of which have crystals - are at the middle of the chain and the newest are at the chain periphery.

We further demonstrated that the interaction between McaA and McaB is correlated to changes in MamK dynamics. When the McaA-McaB interaction is removed through protein truncation or localization changes, MamK dynamics shift such that there are more cells exhibiting the moving-bleach-spot dynamics. This led to hypothesize that MamK dynamics locally controlled based on where McaAB interactions occur. McaB is at the crystal-containing magnetosomes in WT AMB-under norma biomineralization conditions while McaA localizes from cell pole-to-pole along the positive curvature in a dashed pattern in all mutants and growth conditions tested (10). Previous co-localization studies of McaA-Halo and McaB-GFP in WT AMB-showed that McaB-GFP tends to localize within the gaps of the McaA-Halo signal (10). The inverted localization pattern suggests that McaA and McaB only interact at the interface of their territories along the magnetosome chain. Alternatively, the inverted localization pattern could be a result of dynamic McaA-McaB-MamK activity that results in minimized McaA-McaB interactions. In either case, there is a confinement of McaA-McaB interactions, which in turn generates localized zones of MamK regulation.

Altogether, we propose the following model for magnetosome chain organization in AMB-1 (Fig. 7B). McaB is recruited to a magnetosome that has begun biomineralization. This increases the proximity of McaB to the positive curvature membrane-localized McaA and MamK filaments at the magnetosome chain, allowing for protein-protein interactions. The interactions between the three proteins activate McaA and McaB to alter MamK filament directionality locally, which consequently moves the attached magnetosomes. The shifting of these crystal-containing magnetosomes causes some to join into a subchain and others to dislodge from an existing subchain. This in turn opens up space between crystal-containing magnetosomes for new, empty magnetosomes to form. The resulting magnetosome chain therefore has subchains of crystal-containing magnetosomes that are interrupted by empty magnetosomes.

Several pieces of this model remain to be resolved. Firstly, are there other proteins involved in patterning magnetosomes in AMB-1? Though we have presented evidence that McaA and McaB directly interact with MamK, we could not prove a direct interaction between McaA and McaB *in vitro* This leaves the possibility that there are other protein contributors to magnetosome organization not identified here. Secondly, we still do not know how MamK is altered by McaAB at the biochemical level. Possibilities include changes in filament polarity, sites of polymerization, or ATPase activity. Lastly, our model relies on McaAB interactions to specifically occur at the crystal-containing magnetosomes, which we have not demonstrated here. Future work should aim to address these unknowns.

### Changing MamK dynamics as a strategy to diversify magnetosome chain organization

Across MTB, we find diverse types of magnetosome chains. For example, there are species with multiple chains that are parallel, or strikingly, perpendicular to each other (33, 34). How did different types of magnetosome chain arrangements emerge across MTB species? One way is to change the machinery that controls magnetosome positioning. In *Desulfovibrio magneticus* RS-1, for example, second bacterial actin Mad28 and a suite of coiled-coil proteins are required for magnetosome chain assembly and organization in addition to MamK (11). Our work here reveals a possible second strategy where MamK is regulated to produce different magnetosome chains. This serves as a possible explanation for the difference in the magnetosome chain patterning between AMB-1 and its close relative *M. gryphiswaldense* MSR-1 (MSR-1). Both organisms MamK, MamY, and MamJ to build magnetosome chain, yet WT MSR-1 has magnetosome chain with all crystal-containing magnetosomes at the chain center, just like AMB- Δ*mcaAB* (9, 17). Furthermore, in MSR-1, MamK polymerization has been reported to occur at the cell poles, thus pushing the MamK filaments towards midcell, resulting in a moving-bleach-spot dynamic when observed with FRAP (17). The ability to produce a singular crystal chain is not a unique property of MSR- MamK since MSR- Δ*mamK* expressing AMB-1 *mamK* produces a similar magnetosome chain observed by TEM (35). Collectively, it seems likely that the difference in magnetosome chain patterning between AMB-1 and MSR-1 is due to different MamK behavior mediated by McaA and McaB. This case might be one example of a broader mechanism underlying organization diversification of magnetosomes and other subcellular structures in bacteria

### McaA and McaB - bacterial actin binding proteins?

Eukaryotic cells are known to have many actin binding proteins that regulate all aspects of actin’s properties, ranging from capping proteins that prevent further actin polymerization to cross-linking proteins that stabilize higher order filament structures (36). The massive collection of actin binding proteins allows actin to be a multi-functional filament that can have different architectures and dynamics depending on the subset of actin binding proteins that it is interacting with. Compared to the impressive repertoire of eukaryotic actin binding proteins characterized, the discovery of analogous proteins in bacterial systems have lagged behind (37). One aspect is the lack of conservation between eukaryotic and bacterial actin binding proteins, which is largely driven by the divergence of actin filament structure across bacterial actins (37, 38), hindering the computational identification of potential regulators. McaA and McaB interaction with MamK, and MamK’s behavioral change in their absence, opens the possibility that McaA and McaB are bacterial actin binding proteins. Further studies on the biochemical mechanism of McaAB influence on MamK behavior would not only reveal whether they are true actin binding proteins, but also serve as an example that will guide discoveries to other bacterial actin regulators.

## MATERIALS AND METHODS

### Bacterial strains and growth

*Magnetospirillum magneticum* sp. AMB-1 was cultured as previously described (10). Briefly, AMB-1 was grown on MG medium supplemented with 30µM ferric malate and 1x Wolfe’s vitamin solution. Solid medium is prepared by adding 0.7% agar before autoclaving. AMB-1 colonies grown o MG plates were used to inoculate 1.5mL liquid MG in 1.5mL microcentrifuge tubes, then grown at 30°C for 2-3 days. These stock cultures were maintained at room temperature for a maximum duration of 2 weeks and diluted 100-fold into fresh MG medium for growth and C_mag_ measurements, TEM, and fluorescence microscopy. AMB-1 growth for co-IP experiments are detailed in a different section. Unless indicated otherwise, AMB- was cultured in glass culture tubes at 30°C in 10% oxygen without shaking for 1-2 days in a microaerobic glove box. If anaerobic medium was used, the medium was prepared by bubbling MG with N_2_ gas for at least 10 minutes before autoclaving in sealed Balch tubes. Ferric malate, Wolfe’s vitamin solution, and inoculum were introduced through a sterile needle and syringe. When needed, kanamycin was added for a final concentration of 7µg/mL and 10µg/mL for liquid and solid medium, respectively.

OD and coefficient of magnetism (C_mag_) readings for AMB-1 were taken on a Spectronic 20D+ at 400nm. C_mag_ determined by taking OD_400_ readings with stirbar magnet held parallel and perpendicular to the sample. C_mag_ was calculated by taking the ratio of the two OD_400_ readings. A C_mag_ of 1 indicates no magnetic alignment.

*Escherichia coli* DH5ɑ, WM3064, and BL21 were grown in standard LB medium at 37°C, with 50µg/mL kanamycin, 25µg/mL chloramphenicol, 100µg/mL carbenicillin, and/or 5µM diaminopimelic acid (DAP) when needed, unless stated otherwise. Growth of *E coli* strains used for bacterial two hybrids are explained in a different section.

### Non-biomineralization conditions

Al non-biomineralization cultures were grown in MG medium without the ferric malate solution in glassware treated overnight with 30mM oxalic acid then rinsed with MilliQ water three times to remove residual iron. To reduce existing magnetite n AMB-cell, AMB-1 cultures were passaged in 10mL non-biomineralization MG in acid-washed tubes three times, where each passage received a 100µL inoculum of the previous passage and incubated for 2 days microaerobically at 30°C. The final culture was inoculated at a 1:100 dilution and grown at 30°C with constant stirring in the microaerobic glove box to incorporate 10% oxygen to further inhibit biomineralization.

### Genetic Manipulation

All strains and plasmids used in this study be found in the Supplementary Materials. Plasmids used to express genes in AMB-1 or *E. coli* were cloned in *E. coli* DH5ɑ as previously described (10), with a few adjustments. Oligonucleotides were purchased from Integrated DNA technologies and PCR was done using Q5 High Fidelity DNA Polymerase (New England Biolabs). Constructed plasmids were sequence verified through the UC Berkeley DNA sequencing facility or Plasmidsaurus (www.plasmidsaurus.com). Briefly, to generate deletions in AMB-1, a suicide plasmid carrying two homologous regions flanking the target gene was transformed into *E coli* WM3064. The plasmid is introduced into AMB-1 through conjugation, then kanamycin and 2% sucrose are used to select for the first and second recombination events, respectively. Deletions are PCR amplified and sequence verified.

### Cell lysis and *in vivo* co-immunoprecipitation

2L glass bottles with 2L of MG were inoculated with 2 10mL seed cultures of AMB-grown microaerobically for 2 days. These 2L cultures incubated for 2 days at 30°C in 10% oxygen without shaking. If applicable, 10mL of the culture was taken to determine C_mag_ and perform microscopy. To harvest, cells were centrifuged in 500mL centrifuge bottles at 11,000 g with a JA-10 rotor and Beckman J2-21M centrifuge or GS-3 rotor and Sorvall RC-5B centrifuge. Cells were resuspended in 3-5mL medium and transferred to a 15mL falcon tube. The residual medium removed through final centrifugation at 8000 g for 6min. Cell pellet recorded, flash frozen in liquid nitrogen, and stored at -80°C until se AMB-1 was subject to chemical lysis as previously reported (39). Briefly, cell pellets were resuspended in 1mL Lysis Buffer A (10mM Tris-HCl, 150mM NaCl, 0.5mM EDTA, 0.5% NP-40, pH 7.5 supplemented with 2µg/mL pepstatin A, 2µg/mL leupeptin, 2mM PMSF, and 0.5mg/mL lysozyme) and incubated at RT for 15min without rotation. 3mL of Lysis Buffer B (20mM HEPES-KOH, 50mM NaCl, 1.25mM CaC _2_ pH 7.5 with 2mM DTT and 5µg/mL DNase I) was added and samples were incubated at 4°C while rotating end-to-end for 45min. Samples were then centrifuged at 14,600 x g for 30min at 4°C. The supernatant was collected into a new 15mL Falcon tube and either used for co-IP immediately or flash frozen in liquid nitrogen and stored at -80°C until use.

For co-IPs to be evaluated by Western blot, the volume of lysate used was normalized according to the cell pellet mass harvested, then diluted two-fold with Wash Buffer (10mM Tris-HCl, 150mM NaCl, pH 7.4). 20µL of GFP-Trap agarose beads (ChromoTek) were equilibrated three times by resuspending in 500µL Wash Buffer followed by centrifugation. All centrifugation steps here were carried out at 2500 x g at 4°C, unless indicated otherwise. The diluted lysate was applied to the equilibrated beads in a 15mL Falcon tube and rotated end-to-end for 1hr at 4°C. The solution was centrifuged for 6min. The beads were transferred to a 1.5mL microcentrifuge tube and washed three times with 500µL Wash Buffer, centrifuged for 3min between each wash. To elute, the beads were resuspended in 100µL of 2x Laemmli sample buffer and incubated at 10min at 95°C. The samples were centrifuged for 3min and the supernatant was collected as the elution.

The co-IP protocol adjusted for samples to be sent for spectrometry. 10 2L cultures were pooled (20L total). Cell lysis was done with 5mL Lysis Buffer A and 15mL Lysis Buffer B in a 50mL Falcon tube. The lysate was clarified by being split evenly into 6 15mL falcon tubes before centrifugation to minimize debris contamination. 30µL of the GFP-Trap agarose bead slurry was used for the co-IP. Elution was done by resuspending the samples in 50µL of 200mM glycine, pH 2.5 and vigorously pipetting up and down for 30s. The sample was centrifuged at 2500 g for 3min at 4°C The supernatant was transferred to a new tube and neutralized with 5µL of 1M Tris, pH 10.4. The beads were then eluted a second time to increase protein abundance.

### Mass spectrometry

Proteins were precipitated using trichloro acetic acid (TCA) precipitation. 100% TCA was added to the co-IP elution for a final TCA concentration of 20%. Samples were ncubated o ice for 1hr, then centrifuged at 16,000 x g for 10min at 4°C. The protein pellet was washed three times with cold 100µL 0.01M HCL in 90% acetone, then allowed to dry at RT. Protein pellet mass was measured before proceeding. Precipitated protein samples were resuspended in 80µL 100mM Tris-HCl, 8M urea, pH 8.5. 100mM TCEP was added for a final concentration of 5mM, then incubated at RT for 20min. Fresh 500mM iodoacetamide was added for a 10mM final concentration, then incubated at RT for 15min in the dark. Samples were diluted 4-fold using 100mM Tris-HCl, pH 8.5. Filter-sterilized 100mM CaCl_2_ was added for a final concentration of 1mM before adding 1µL of 0.5µg/µL sequence-grade modified trypsin (Promega). Samples were incubated overnight at 37°C in the dark, then formic acid was added for a 5% final concentration. Samples were then desalted using C18 Spec tips (Agilent). The Spec tip was washed with 200µL HPLC-grade MeOH, followed by three washes with 20µL 5% acetonitrile/5% formic acid. Sample was pushed through the tip, washed three times with 200µL 5% acetonitrile/5% formic acid, eluted with 2 passes of 100µL 80% acetonitrile/5% formic acid, then dried.

Mass spectrometry was performed at the Proteomics/Mass Spectrometry Laboratory at University of California, Berkeley. A nano LC column was packed in a 100-µm inner diameter glass capillary with an integrated pulled emitter tip. The column consisted of 10cm of Polaris c18 5-µm packing material (Varian). The column was loaded and conditioned using a pressure bomb. The column then coupled to electrospray ionization mounted Thermo-Fisher LTQ XL linear ion trap mass spectrometer. An Agilent 1200 HPLC equipped with a split line so as to deliver a flow rate of 1µl/min was used for chromatography. Peptides were eluted with a 90-minute gradient from 100% buffer A to 60% buffer B. Buffer A was 5% acetonitrile/0.02% heptafluorobutyric acid (HBFA); buffer B was 80% acetonitrile/0.02% HBFA. Collision-induced dissociation and electron transfer dissociation spectra collected for each *m/z* Protein identification and quantification and analysis done with Integrated Proteomics Pipeline-IP2 (Bruker Scientific LLC, Billerica, MA, http://www.bruker.com)) using ProLuCID/Sequest (40), DTASelect2 (41, 42), and Census (43, 44). Spectrum raw files were extracted into ms1 and ms2 files from raw files using RawExtract 1.9.9 (http://fields.scripps.edu/downloads.php) 10, and the tandem mass spectra were searched against MIT database (42, 45). Proteins encoded in AMB-1 magnetosome islet (MIS) downloaded from NCBI on Sept. 1, 2022 and included for analysis.

### Western Blotting

Western blots were done with protein samples separated via SDS-PAGE using Mini-PROTEAN TGX Stain-Free (BioRad). Prior to transfer, gels were imaged for total protein content using Stain-Free imaging. The primary antibodies and their dilutions used in this study are as follows: anti-GFP (Abcam ab 6556, 1:2500 and Invitrogen GF28R, 1:2500), anti-Halotag (Promega G921A, 1:1000), and anti-Flag (Sigma F3165, 1:2500). The secondary antibodies and their dilutions are as follows: anti-mouse goat HRP conjugate (Invitrogen A24512, 1:5000), anti-rabbit goat HRP conjugate (Biorad, 170-5046, 1:10,000). Blots were developed using Western Lighting Plus-ECL (PerkinElmer) and imaged with the BioRad ChemiDoc MP Imaging System. All full-sized blots be found n the Source Data file.

### Epifluorescence microscopy

10mL of AMB-1 microaerobically grown for 2 days at 30°C until mid-to-late exponential phase (OD_400_ of 0.200-0.280). 1-1.5mL of this culture was centrifuged for 4min at 8000 x g. The pellet was resuspended in 10µL of residual medium and 0.7µL was applied on a glass slide Imaging was done on a Zeiss Axio Observer Inverted microscope with Zen software (Zeiss), and images were handled using Fiji. To score images for localization, images were randomized using the Cell Counter plugin.

### Fluorescence recovery after photobleaching

10mL AMB-cultures were grown microaerobically until OD_400_ reached ∼0.1. The cultures were concentrated to 10µL, of which 0.7µL was applied between the bottom of a glass bottom dish (MatTek) and a 450µL 2% agarose gumdrop that minimizes cell movement. Cells imaged inverted Carl Zeiss LSM880 FCS laser scanning confocal microscope with an objective lens Plan-Apochromat 100x/1.40 oil DIC at RT. Cells were imaged with a 488nm laser at 3% power every 30s for 15min with Definite Focus autofocusing (Zeiss). At t=1min, a small region of the cell was photobleached with the 488nm laser at 100% power. Images were acquired through LSM880 Zen software (Zeiss). Fiji was used to analyze the images and generate kymographs.

### Transmission electron microscopy

AMB-1 cells to be used for TEM were grown in 10mL cultures microaerobically for 2 days at 30°C. 1-1.5mL of the culture concentrated to 10µL and applied to Formvar/Carbon 300 Mesh copper grid (Electron Microscopy Sciences) that have been glow discharged with Pelco Easiglow. After 5min incubation at RT, excess cells were removed by washing the grid three times in MilliQ water. Cells were imaged on an FEI Tecnai 12 transmission electron microscope with an accelerating voltage of 120 kV. Images were captured using a Rio 16 4K CMOS camera and Gatan Digital Micrograph software Grids carrying cells to be used for scoring randomized before imaging and scoring.

### Protein purification

To purify McaA and McaB, the predicted cytoplasmic side of both proteins were tagged, then expressed in BL2 *E coli* McaA (aa400-776) was N-terminally tagged with a Strep-tag and McaB (aa 28-219) was N-terminally tagged with maltose binding protein (MBP). MBP was also expressed from BL21. BL21 harboring the recombinant protein expression plasmids were inoculated as 1:100 dilution from overnight seed culture grown in 2xYT without inducers and grown at 37°C shaking at 200rpm. When OD_600_ reached ∼0.5, Strep-McaA and MBP expression was induced with 0.1mM IPTG and MBP-McaB expression was induced with 0.5mM IPTG. Cultures were incubated for another 3-4hrs at 37°C shaking at 200rpm. Cells were harvested at 10,800 x g for 20min at 4°C, flash frozen with liquid nitrogen, and stored at -80°C until use To lyse, cell pellets were resuspended in 35mL 10mM Tris-HCl, 150mM NaCl, pH7.4 supplemented with 1µg/ml pepstatin A, 2µg/ml leupeptin, and 1mM PMSF, and lysed with a french press at 18,000 PSI. Lysate was clarified 14,600 x g for 30min at 4°C, and stored at 4°C overnight. Lysates were filtered through a 0.22µm PES syringe filter and applied to either a StrepTrap HP (GE Healthcare) or MBPTrap HP (Cytiva) column using an ӒKTA pure FPLC system with 5ml/min flow rate. Samples were washed with 20 column volumes (CV) with 10mM Tris-HCL, 150mM NaCl, pH 7.4 and eluted with 5CV with either 10mM Tris-HCl, 150mM NaCl, 2.5mM desthiobiotin, pH 7.4 for StrepTrap or 10mM Tris-HCl, 150mM NaCl, 10mM maltose, pH 7.4 for MBPTrap. Fractions were evaluated with SDS-PAGE and those containing the protein of interest were pooled and concentrated with Amicron Ultra 10K centrifugal filter units. Samples were further purified via size exclusion chromatography using a HiLoad 16/600 Superdex 75 pg column (GE Healthcare) with 10mM Tris-HCL, 150mM NaCl, pH 7.4. Fractions with pure proteins were flash frozen and stored at -80°C until use. Protein concentrations were determined using the Micro BCA Protein Assay Kit.

Untagged recombinant MamK purified from *E. coli* using ammonium sulfate precipitation previously described (20), with a few adjustments. After the second 20% ammonium sulfate cut, MamK samples are dialyzed into depolymerization buffer (10mM CAPs, 10mM EDTA, 2mM DTT, pH 9.6) at 4°C for approximately 8h. The sample was further purified using anion exchange chromatography (DEAE TOYOPEARL, 2.3cm × 12cm, Tosoh, Japan) and gel filtration chromatography (Superdex 200 Increase 10/300 GL, Cytiva, USA) with depolymerization buffer. The monomeric MamK fractions were pooled, and aliquots were stored at −80°C. For experimental use, purified MamK is exchanged into polymerization buffer (10mM Tris-HCl, 25mM KCl, 2mM MgCl_2_ 1mM DTT, pH 7.5) using a desalting spin column (Zaba Spin Desalting Column, Thermo Fisher, USA).

### Pelleting Assay

Protein samples were diluted to 1µM in polymerization buffer (10mM Tris-HCl pH 7.5, 25mM KCl, 2mM MgCl_2_ 1mM DTT). Proteins in mixtures each had a concentration of 1µM. Final concentration of 1mM ATP was added if needed Samples were ncubated at 25°C for 10min, then centrifuged at 100,000 g for 30min at 4°C. The supernatant and pellet fractions were recovered and analyzed by SDS-PAGE.

### *In vitro* pull down

10µg of each protein was mixed into a total of 500µL of equilibration buffer (10mM Tris-HCl, 150mM NaCl, pH 7.4) and incubated at 4°C for 1hr with end-to-end mixing. All following steps were carried out at 4°C. For negative controls, 10µg of each protein were incubated separately. 20µL Strep-tactin resin (IBA Lifesciences) was equilibrated by washing three times with 100µL equilibration buffer and centrifuging at 2500 g for 3min at 4°C between each wash. Protein samples were centrifuged at 16,000 x g for 5min to remove any aggregates. The supernatant was added to the equilibrated Strep-tactin resin in a 1.5mL microcentrifuge tube and incubated for 1hr while mixing end-to-end. The sample was centrifuged at 2500 g for 3min, and the flowthrough supernatant was removed. The resin was washed three times using 500µL equilibration buffer, then eluted with 50µL 10mM Tris-HCl, 150mM NaCl, 2.5mM desthiobiotin, pH 7.4 by pipetting vigorously for 30s The elution was collected by centrifuging for 3min at 2500 x g. Samples were evaluated with SDS-PAGE and Coomassie staining.

### Bacterial adenylate cyclase two hybrids assay (B2H)

*E. coli* DHM were transformed with two plasmids, each carrying either the T18 or T25 fusion and incubated at 30°C overnight with 50µg/mL kanamycin and 100µg/mL carbenicillin for selection. A single colony was used to inoculate 80µL LB with kanamycin, carbenicillin, and 0.5mM IPTG in a 96 well plate, and incubated at 30°C overnight shaking at 200rpm. 3µL of the culture spotted onto M63 agar supplemented with 25µg/mL kanamycin, 50µg/mL carbenicillin, 0.5mM IPTG, and 40µg/mL X-gal and incubated at 30°C for 5-8 days before imaging with an iPhone 15 camera.

### Multiple sequence alignment

McaA homologs were first identified through a BlastP search of AMB-1 McaA Dec. 27, 2024. Homolog sequences with expectation value below 1e-5 from this initial search were downloaded from NCBI on the same day. The homolog sequences were then aligned on Jan. 14, 2025 using MAFFT (Version 7.511) using the iterative refinement algorithm L-INS-i (46). The alignment was visualized using Jalview (version 2.11.4.1) (47).

### McaA and McaB protein sequence analysis

McaA protein structure was predicted using AlphaFold3 on the Alphafold Server (https://alphafoldserver.com/) o Nov. 12, 2022 (30). Deepcoil2 (https://toolkit.tuebingen.mpg.de/tools/deepcoil2) was used to predict coiled-coi domains, “a” position amino acids, and “d” position amino acids of McaB (29). The predicted domains and amino acids were mapped onto predicted McaB structures generated by AlphaFold3 on Oct. 20, 2025. Protein structures were visualized using UCSF ChimeraX (48).

## Data Availability

The data from the IP-MS and FRAP experiments can be found in Supplementary Materials.

## Supporting information

Source Data 2: IPMS Experiments

Source Data 3: FRAP Experiments

Source Data 4: Gels

Source Data 5: Strains and Plasmids

Source Data 6: Statistics

## ACKNOWLEDGEMENTS

We thank Lor Kohlstaedt and Robert Maxwell at the Vincent J Coates Proteomics/Mass Spectrometry Laboratory at UC Berkeley for performing the mass spectrometry for this study. We thank Denise Schichnes, Juliana Cho, and Steve Ruzin for their training and support with confocal imaging at the Biological Imaging Facility at UC Berkeley. We thank Danielle Jorgens, Reena Zalpuri, and Misun Kang for training and assistance with TEM at the Electron Microscope Lab at UC Berkeley. A. K. is supported by NIH National Institute of General Medical Sciences (R35GM127114). Y. R. was supported by NSF Graduate Research Fellowship Program (2020254971) and the Berkeley Fellowship for Graduate Study.

## Notes

### Competing Interest Statement

The authors have declared no competing interest.

