## Supplemental Information and Figures for "Magnetosome organelles are organized through interactions between McaA and McaB that alter the dynamics of the bacterial actin-like protein MamK"

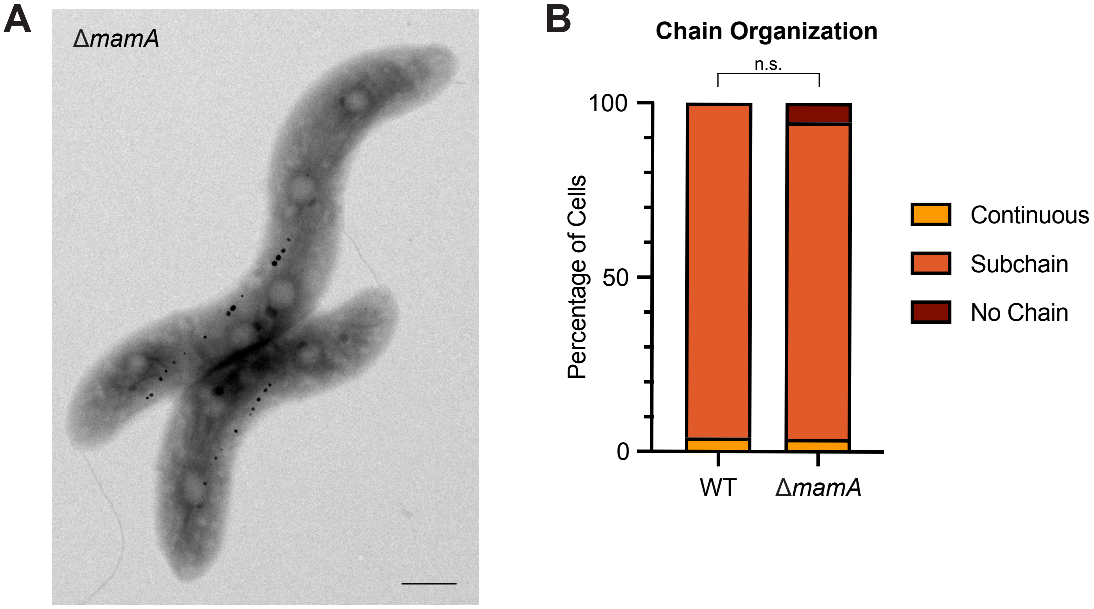


**FIG S1**: MamA is not necessary for crystal subchains. (**A**) Representative TEM image of AMB-1 Δ*mamA* showing the magnetosome chain organization. Scale bar = 0.5µm. (**B**) Quantification of magnetosome chain organization using TEM images. Cells were categorized based on their magnetosome chain phenotype and the y-axis represents the percentage of cells that displayed the indicated chain organization. WT n = 75, Δ*mamA* n = 54. Fisher's exact test was used to determine p-values (n.s. not significant).

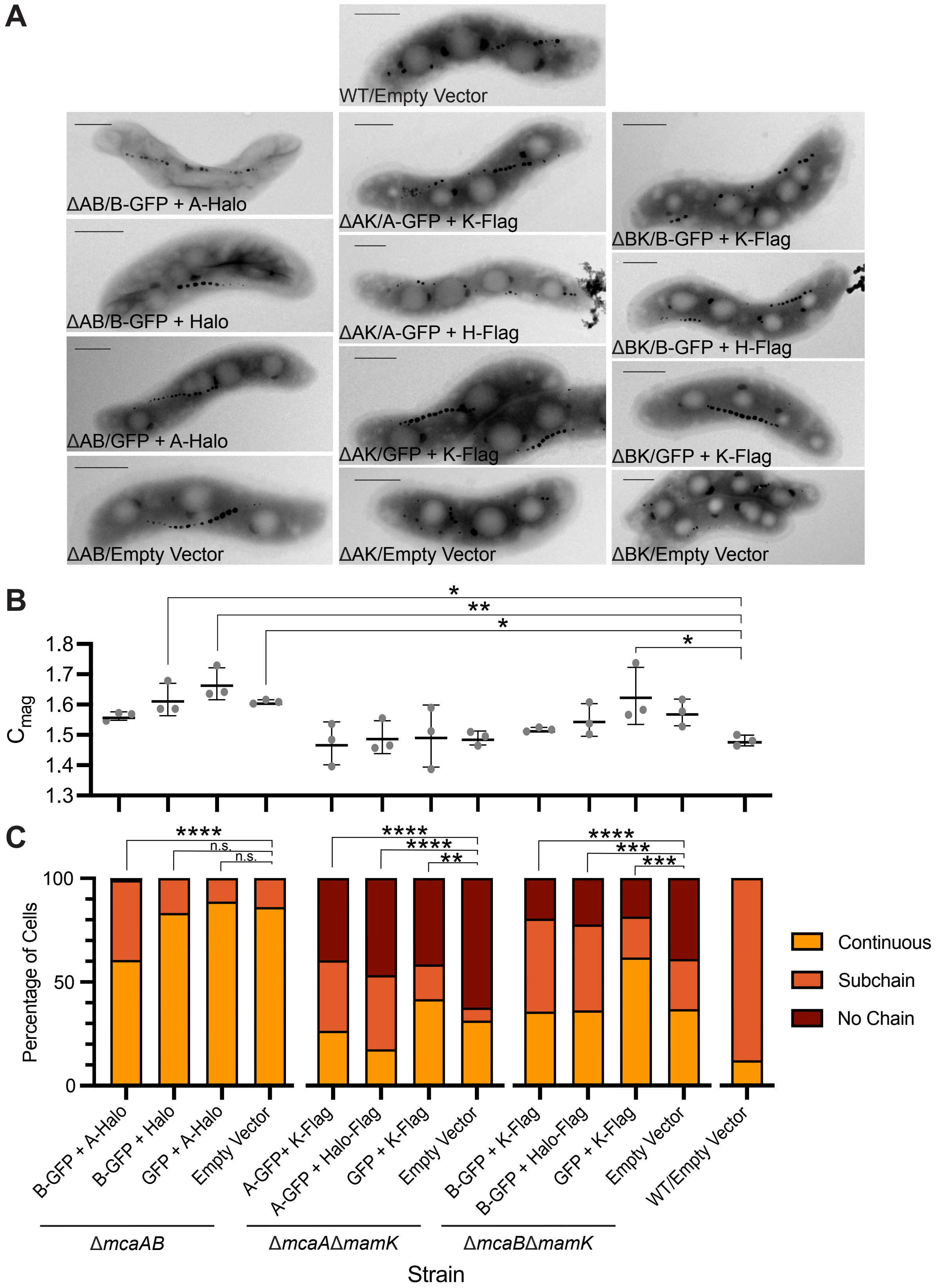
A
WT/Empty Vector
ΔAB/B-GFP + A-Halo
ΔAK/A-GFP + K-Flag
ΔBK/B-GFP + K-Flag
ΔAB/B-GFP + Halo
ΔAK/A-GFP + H-Flag
ΔBK/B-GFP + H-Flag
ΔAB/GFP + A-Halo
ΔAK/GFP + K-Flag
ΔBK/GFP + K-Flag
ΔAB/Empty Vector
ΔAK/Empty Vector
ΔBK/Empty Vector
B
*
**
*
*
1.8
1.7
1.6
1.5
1.4
1.3
C_mag
C
****
n.s.
n.s.
****
****
**
****
***
***
100
50
0
Percentage of Cells
Continuous
Subchain
No Chain
B-GFP + A-Halo
B-GFP + Halo
GFP + A-Halo
Empty Vector
A-GFP+ K-Flag
A-GFP + Halo-Flag
GFP + K-Flag
Empty Vector
B-GFP + K-Flag
B-GFP + Halo-Flag
GFP + K-Flag
Empty Vector
WT/Empty Vector
ΔmcaAB
ΔmcaAΔmamK
ΔmcaBΔmamK
Strain

**FIG S2**: Tagged constructs of McaA, McaB, and MamK can partially complement their respective deletions. (**A**) Representative TEM image of different strains used for co-IPs. Scale bar = 0.5μm. (**B**) The coefficient of magnetism ($C_{mag}$) of the different co-IP strains. The column labels are the same as those in (C). Each point represents one independent culture, and the average and standard deviation is shown. One-way ANOVA with Dunnett's multiple comparisons test was used to compare each strain to both its respective empty vector and WT/empty vector. Comparisons without stars did not have a significant difference. (**C**) Quantification of magnetosome chain organization using TEM images. Cells were categorized based on their magnetosome chain phenotype and the y-axis represents the percentage of cells that displayed the indicated chain organization. n from left to right: 104, 125, 115, 93, 106, 109, 137, 96, 143, 210, 81, 182, 91. Fisher's exact test was used to determine p-values (* $P < 0.05$, ** $P < 0.01$, *** $P < 0.001$, **** $P < 0.0001$, n.s. Not significant). A = *mcaA*, McaA; B = *mcaB*, McaB; K = *mamK*, MamK; H = Halo-tag.

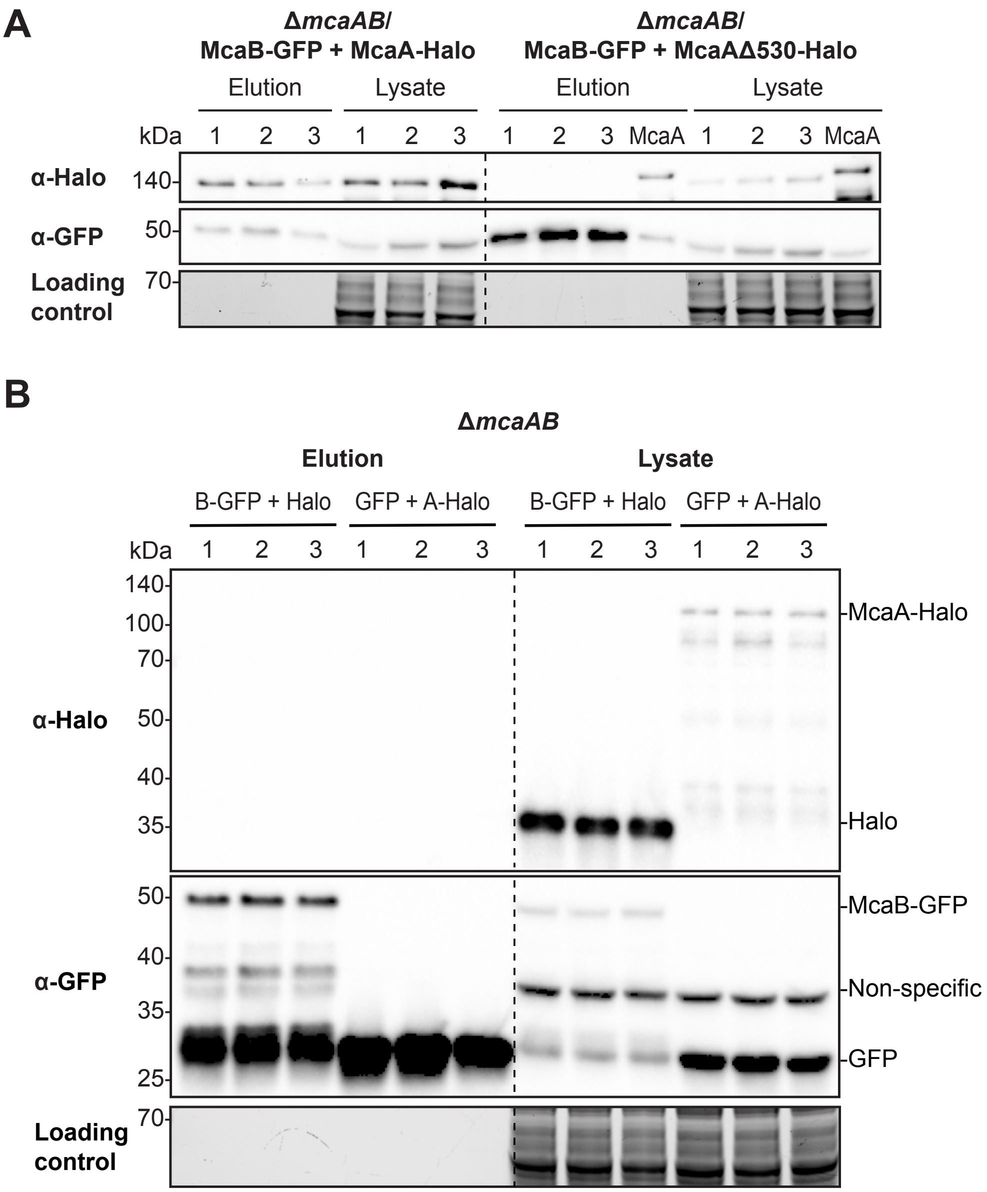


**FIG S3**: McaA and McaB interact with each other *in vivo*. (**A**) Western blots of co-IP elutions and lysates illustrating that McaB-GFP interacts with McaA-Halo, but not McaAΔ530-Halo. Three independent biological replicates are shown. The McaB-GFP and McaAΔ530-Halo co-IPs include a positive control done with full length McaA-Halo (lane titled McaA). (**B**) Western blots of co-IP elutions and lysates with McaB-GFP (B-GFP) and Halo-tag or GFP tag and McaA-Halo (A-Halo) illustrating that the interaction detected between McaB-GFP and McaA-Halo is not due to their tags. All co-IPs were done using the Δ*mcaAB* strain using GFP-Trap agarose beads. Three independent biological replicates are shown. The loading control is a stain-free SDS-PAGE gel.

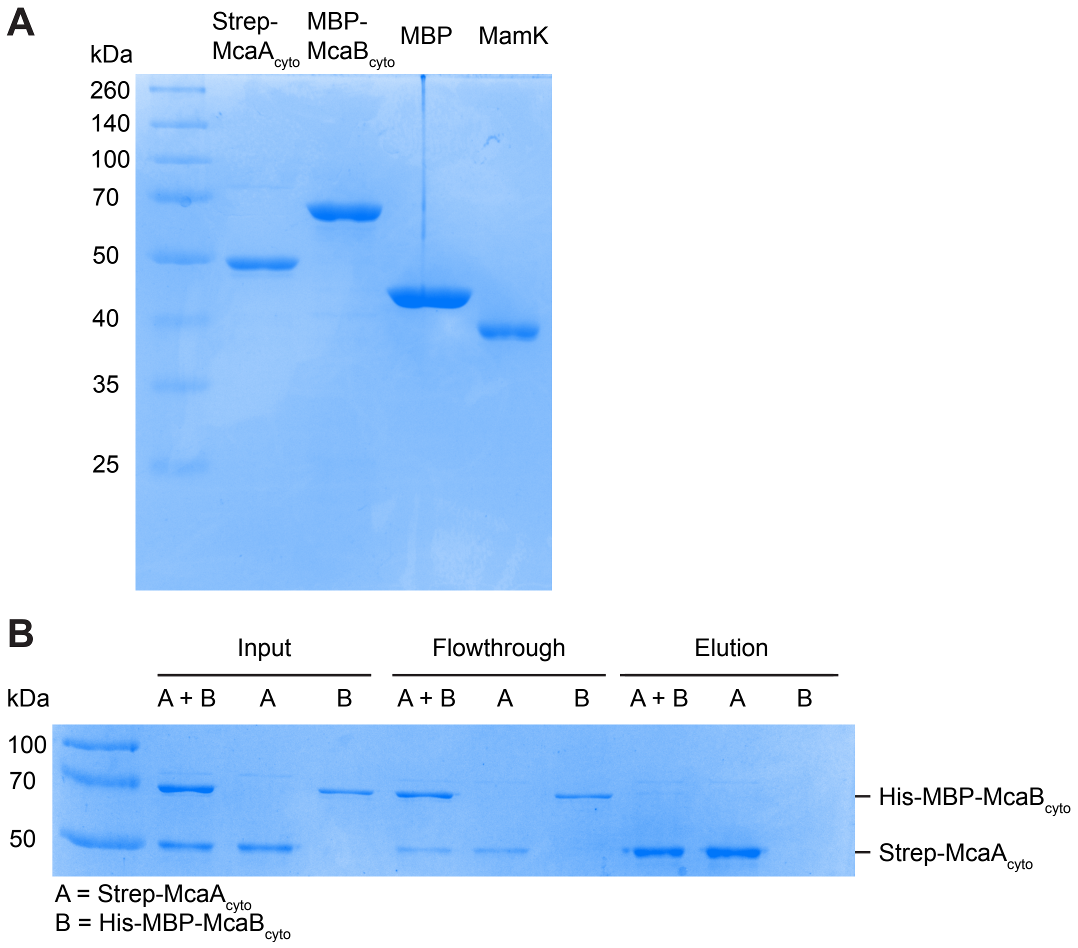


**FIG S4**: Purified cytoplasmic domains of McaA and McaB do not directly interact *in vitro*. (**A**) SDS-PAGE with purified Strep-McaA$_{cyto}$, MBP-McaB$_{cyto}$, MBP, and untagged MamK. (**B**) SDS-PAGE of an *in vitro* pull down with purified Strep-McaA$_{cyto}$ and His-MBP-McaB$_{cyto}$. Strep-McaA$_{cyto}$ and His-MBP-McaB$_{cyto}$ were applied onto Strep-Tactin resin either individually or mixed together (input). Unbound proteins were washed off (flowthrough) and the resin was washed with a buffer. Elution shows that Strep-McaA$_{cyto}$ and His-MBP-McaB$_{cyto}$ do not co-elute. Details of the *in vitro* pull down can be found in the materials and methods. A, Strep-McaA$_{cyto}$; B, His-MBP-McaB$_{cyto}$.

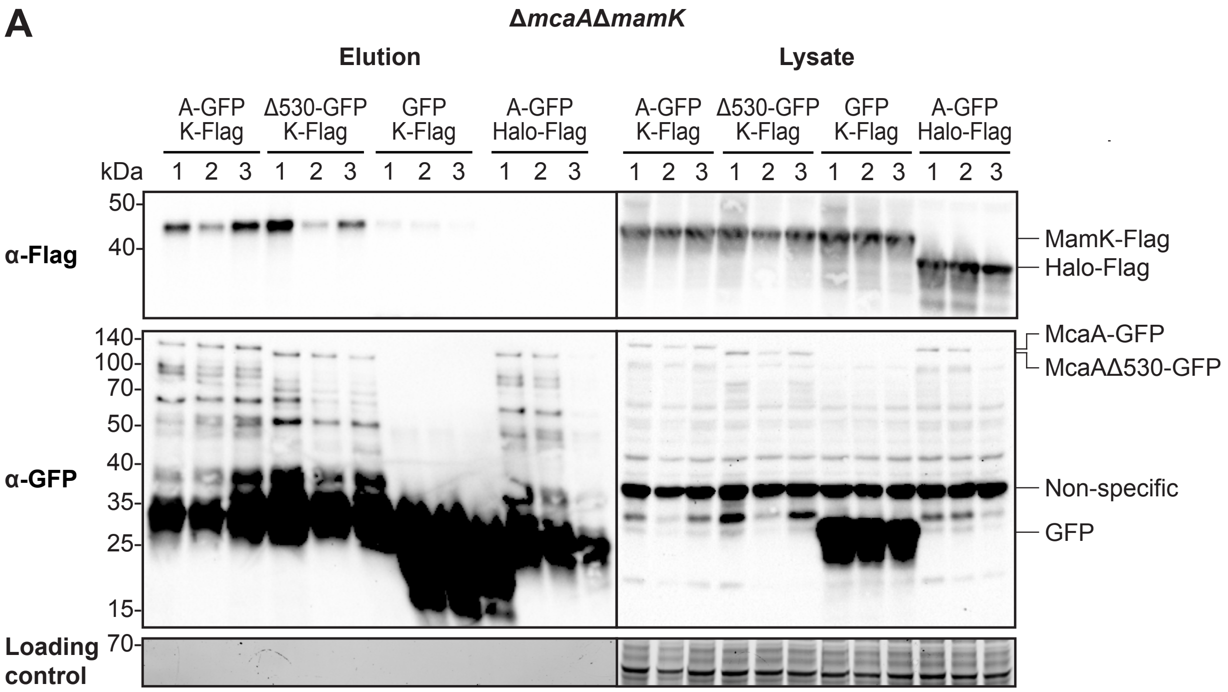


**FIG S5**: MamK interacts with both McaA and McaB *in vivo*. (**A**) Western blots of co-IP elutions and lysates illustrating that MamK-Flag (K-Flag) interacts with both McaA-GFP (A-GFP) and McaAΔ530-GFP (Δ530-GFP). The GFP and Flag tags on McaA and MamK, respectively, play no or minimal role in driving the interaction. (**B**) Western blots of co-IP elutions and lysates illustrating that MamK-Flag (K-Flag) interacts with both McaB-GFP (B-GFP). The GFP and Flag tags on McaB and MamK, respectively, do not drive the interaction. In both (A) and (B), the co-IP was done using GFP-Trap agarose beads and three independent biological replicates are shown. The loading control is a stain-free SDS-PAGE gel.

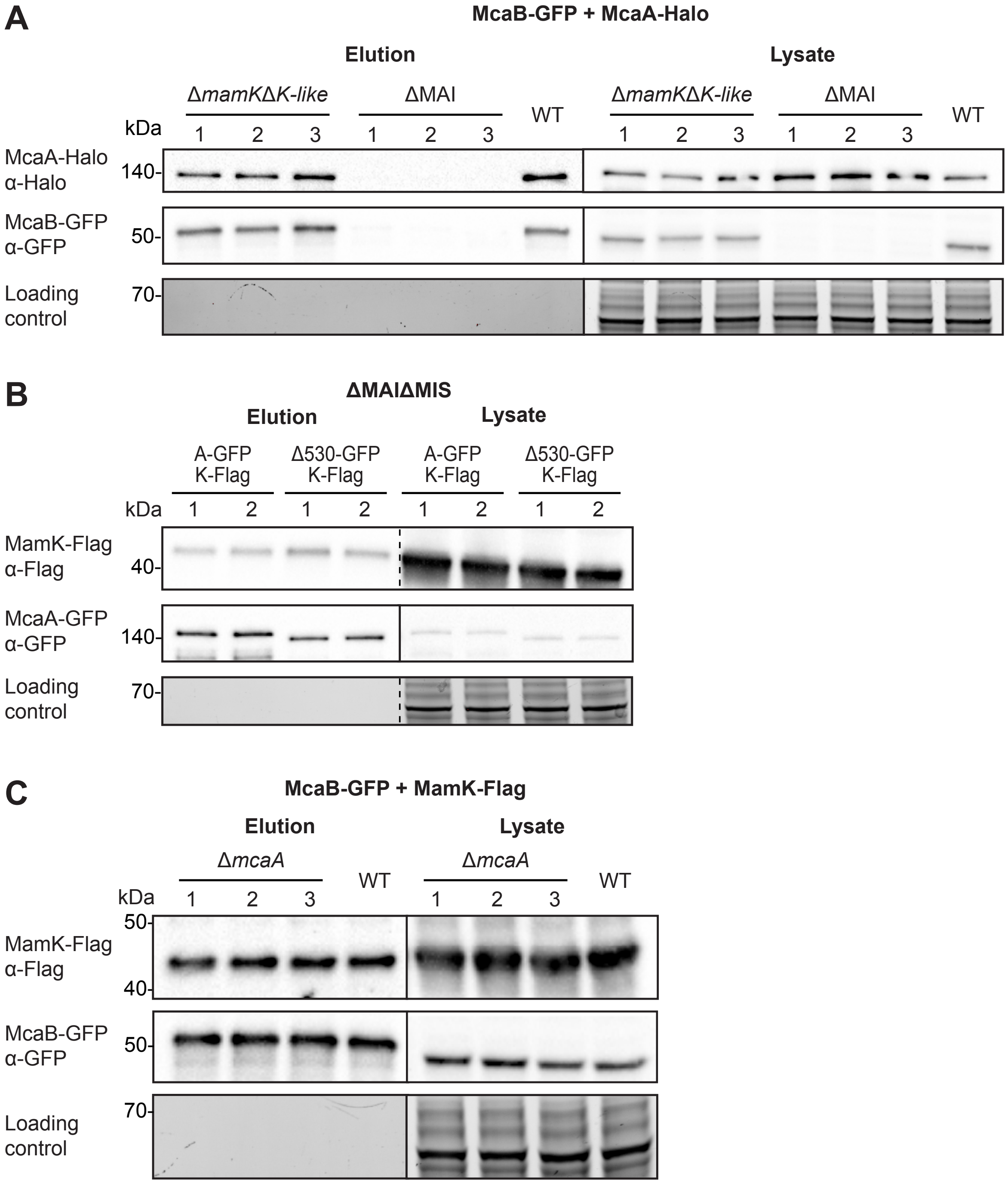


**FIG S6**: McaA, McaB, and MamK form pairwise interactions independent of the third protein. (**A**) Western blots of co-IP elutions and lysates of McaB-GFP and McaA-Halo in Δ*mamK*Δ*mamK-like* and ΔMAI show that the McaA-McaB interaction does not require MamK or MamK-like. McaB-GFP expression is drastically reduced in ΔMAI, evidenced by the lack of McaB-GFP bands in the lysate. WT samples serve as a positive control. (**B**) Western blots of co-IP elutions and lysates of MamK-Flag (K-Flag) with McaA-GFP (A-GFP) or McaAΔ530-GFP (Δ530-GFP) in ΔMAIΔMIS. The results demonstrate that MamK interacts with McaA independent of the 530 region and all other magnetosome proteins. (**C**) Western blots of co-IP elutions and lysates of MamK-Flag with McaB-GFP in Δ*mcaA* show that MamK interacts with McaB independent of McaA. WT samples serve as a positive control. All co-IPs were done using GFP-Trap agarose beads. All blots have independent biological replicates numbered and a stain-free SDS-PAGE gel serves as the loading control.

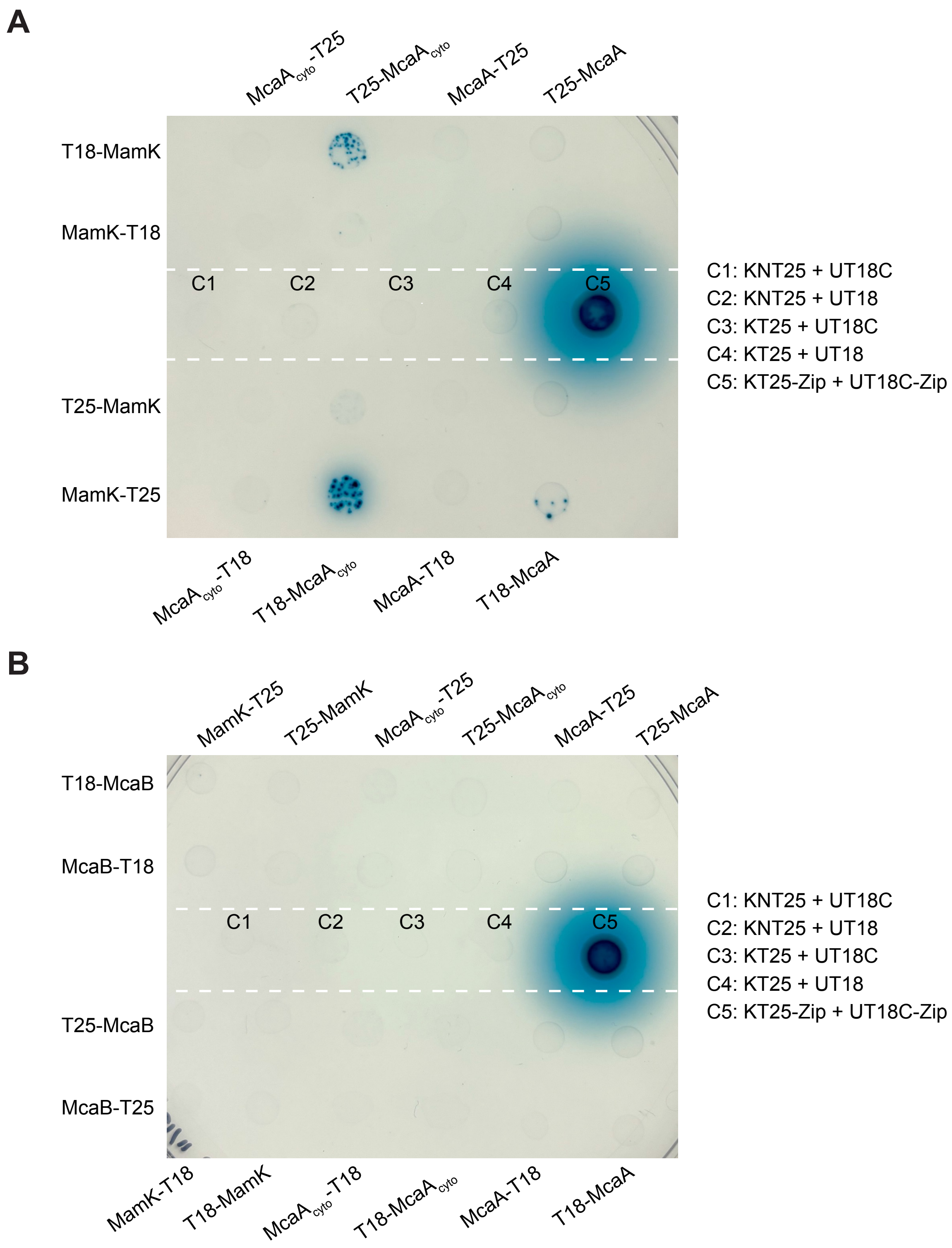


**FIG S7**: Bacterial two-hybrids (B2H) show interaction between MamK and the cytoplasmic side of McaA. (**A**) B2H between MamK and either full length McaA or the cytoplasmic side of McaA (McaA$_{cyto}$) tagged with B2H fragments T18 or T25. The co-expression strains are spotted on M63 agar with IPTG and X-gal. A blue spot indicates an interaction between the two co-expressed proteins. Untagged T18 and T25 negative controls (C1-4) and a leucine zipper positive control (C5) is shown in the center of the plate. (**B**) B2H between McaB and MamK, McaA, and McaA$_{cyto}$. Experimental set up is similar to (A).

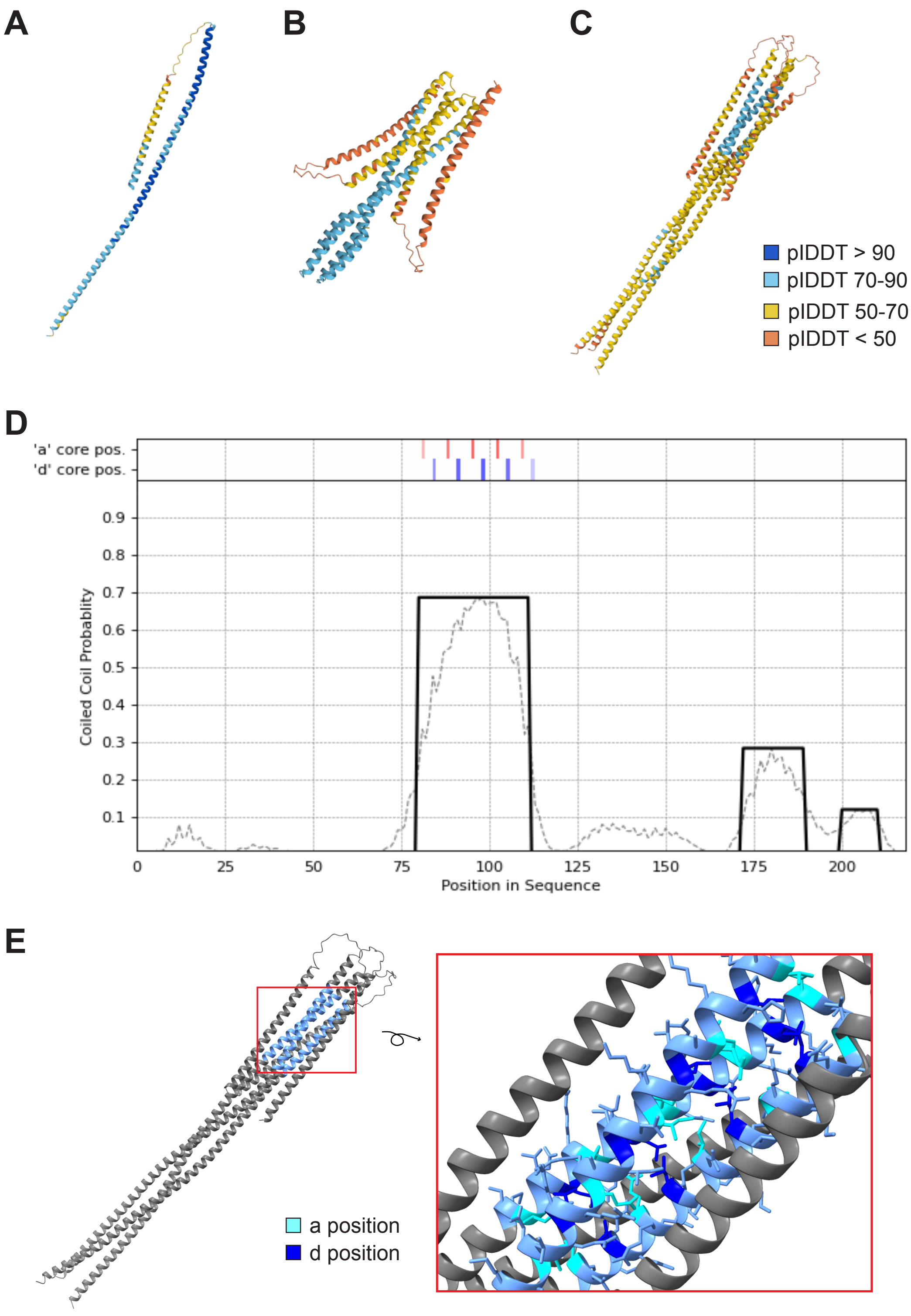


**FIG S8**: McaB is a predicted coiled-coil protein. Alphafold3 predictions of McaB as a monomer (**A**), dimer (**B**), and trimer (**C**). The structures are color-coded based on Alphafold prediction confidence levels (pIDDT), where higher numbers represent greater confidence. (**D**) Probability of coiled-coil domains in McaB determined by DeepCoil2. The x-axis represents the amino acid positions and the y-axis represents the probability of coiled-coils. The red and blue colored bars at the top represent predicted “a” position and “d” position amino acids respectively, where hydrophobic residues occupy in a coiled-coil. (**E**) The predicted coiled-coil domain (light blue) is mapped onto the Alphafold predicted McaB trimer. The inset shows the predicted “a” position (cyan) and “d” position (dark blue) amino acids that form a hydrophobic core typical to coiled-coils.

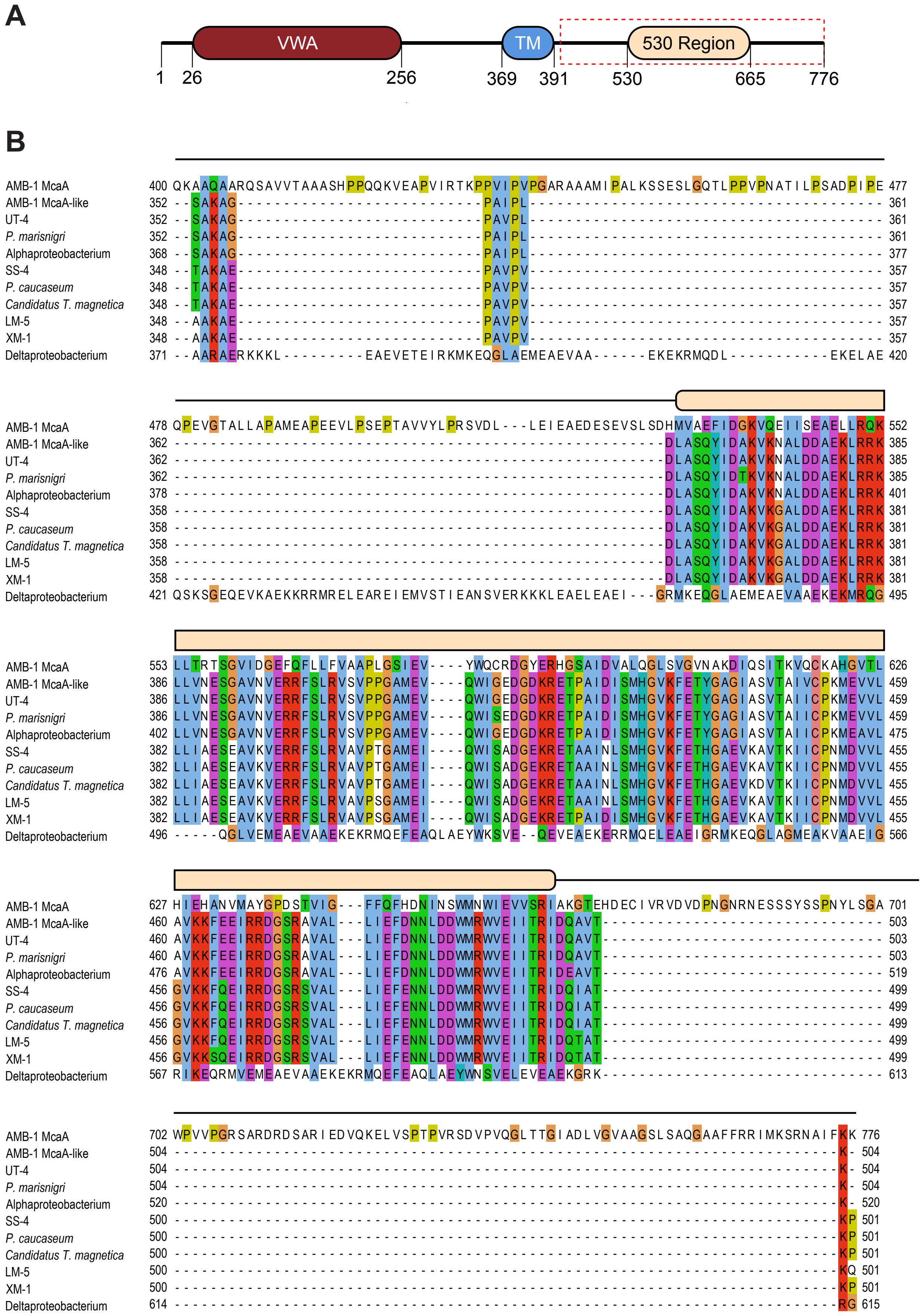

A
VWA
TM
530 Region
1
26
256
369
391
530
665
776
B
AMB-1 McaA
AMB-1 McaA-like
UT-4
P. marisnigri
Alphaproteobacterium
SS-4
P. caucaseum
Candidatus T. magnetica
LM-5
XM-1
Deltaproteobacterium

**FIG S9**: The 530 region of AMB-1's McaA is a conserved region of the cytoplasmic side of McaA. (**A**) Schematic of full length McaA. The amino acid positions are shown. The N-terminus has a von Willebrand factor type A domain (VWA), followed by a transmembrane domain (TM). The cytoplasmic side of McaA is indicated by a red dashed box, which includes the 530 region (amino acids 530-665). (**B**) Amino acid sequence alignment of the cytoplasmic C-terminal side of McaA across different MTB. Both AMB-1 McaA and its homolog McaA-like is included in the alignment. Residues are colored using the Clustal coloring scheme, which assigns colors based on the chemical properties of the amino acid. The schematic from (A) is shown on along the alignment to indicate where the 530 region is. Species name abbreviations: UT-4, *Magnetospirillum* sp. UT-4; *P. marisnigri*, *Paramagnetospirillum marisnigri*; SS-4, *Magnetospirillum* sp. SS-4; *P. caucaseum*, *Paramagnetospirillum caucaseum*; *Candidatus T. magnetica*, *Candidatus Terasakilla magnetica*; LM-5, *Magnetospirillum* sp. LM-5; XM-1, *Magnetospirillum* sp. XM-1.

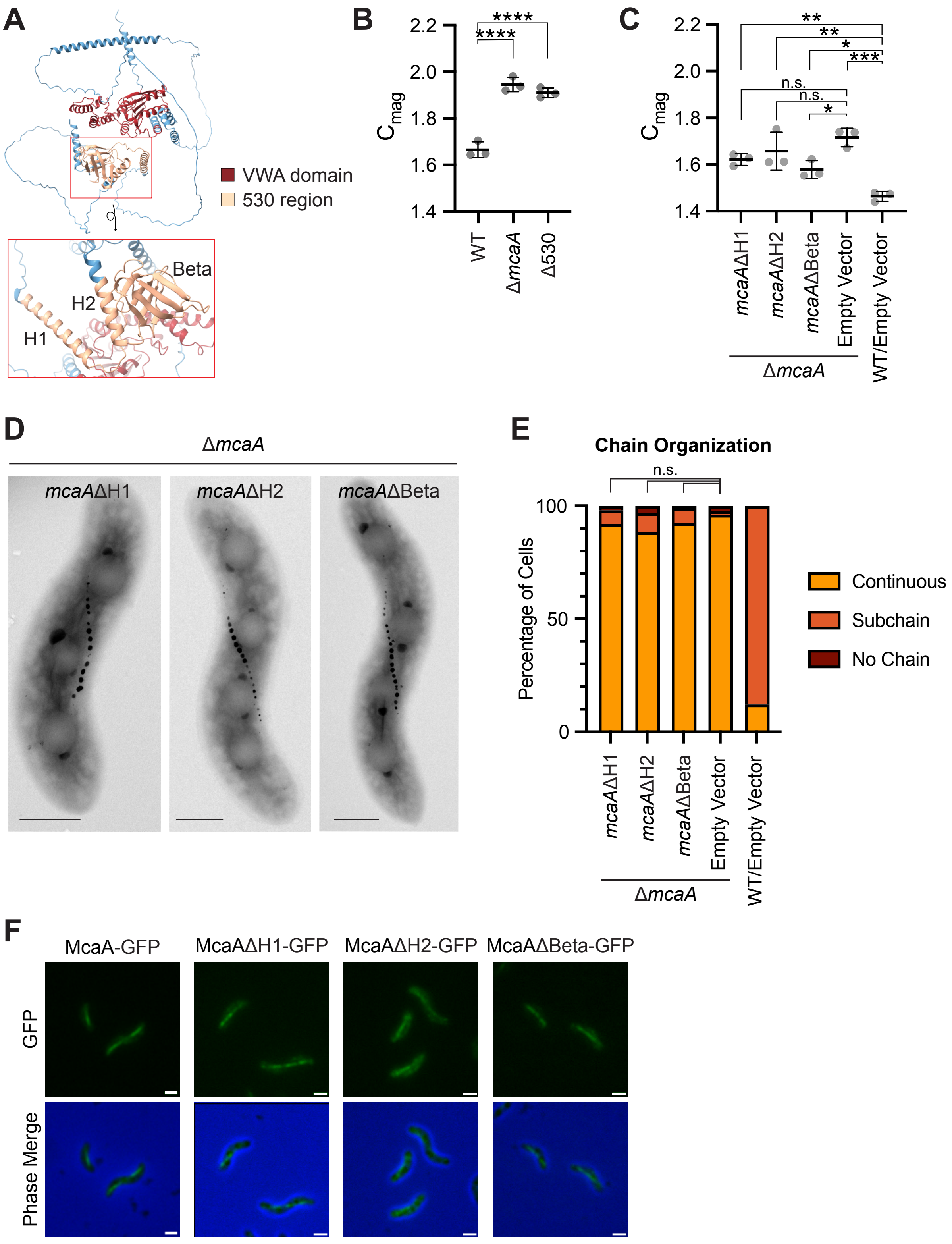

A
VWA domain
530 region
Beta
H2
H1
B
C
Cmag
****
****
WT
ΔmcaA
Δ530
**
**
*
***
n.s.
n.s.
*
mcaAΔH1
mcaAΔH2
mcaAΔBeta
Empty Vector
WT/Empty Vector
ΔmcaA
D
ΔmcaA
mcaAΔH1
mcaAΔH2
mcaAΔBeta
E
Chain Organization
n.s.
Percentage of Cells
Continuous
Subchain
No Chain
F
McaA-GFP
McaAΔH1-GFP
McaAΔH2-GFP
McaAΔBeta-GFP
GFP
Phase Merge

**FIG S10**: The entire 530 region is necessary for crystal subchain formation. (**A**) Predicted protein structure of McaA according to Alphafold 3. The VWA domain and 530 domain are colored red and light yellow, respectively. The inset shows three distinct structural elements of the 530 region: Helix 1 (H1), helix 2 (H2) and a beta sheet (Beta). (**B**) Coefficient of magnetism ($C_{mag}$) measurements of WT, Δ*mcaA*, and Δ530 demonstrate that removing the 530 region increases the $C_{mag}$ to levels similar to that of Δ*mcaA*. (**C**) $C_{mag}$ measurements of strains expressing McaAΔH1-GFP, McaAΔH2-GFP, McaAΔBeta-GFP, or an empty vector in a Δ*mcaA* background and compared to a WT with an empty vector. It is common for WT/empty vector grown with kanamycin to have a lower $C_{mag}$ compared to WT grown without kanamycin. For both (B) and (C), each point represents one independent culture, and the averages and standard deviations are shown. One-way ANOVA with Dunnett's multiple comparisons test was used to determine p-values (* P < 0.05, ** P < 0.01, *** P < 0.001, **** P < 0.0001, n.s. Not significant). (**D**) Representative TEM image of Δ*mcaA* strains expressing McaA with different truncations in the 530 region. All constructs have a C-terminal GFP tag. Scale bar = 0.5μm. (**E**) Quantification of magnetosome chain organization using TEM images indicates that H1, H2, and Beta are all required for McaA function. Cells were categorized based on their magnetosome chain phenotype and the y-axis represents the percentage of cells that displayed the indicated chain organization. n from left to right: 50, 60, 103, 126, 91. Fisher's exact test was used to determine p-values (n.s. Not significant). All groups were significantly different from WT/Empty Vector (P < 0.0001). (**F**) Representative cells expressing GFP-tagged McaA or McaA truncation, showing that each truncated McaA construct is expressed and localized similarly to full-length McaA-GFP. Scale bar = 1μm.

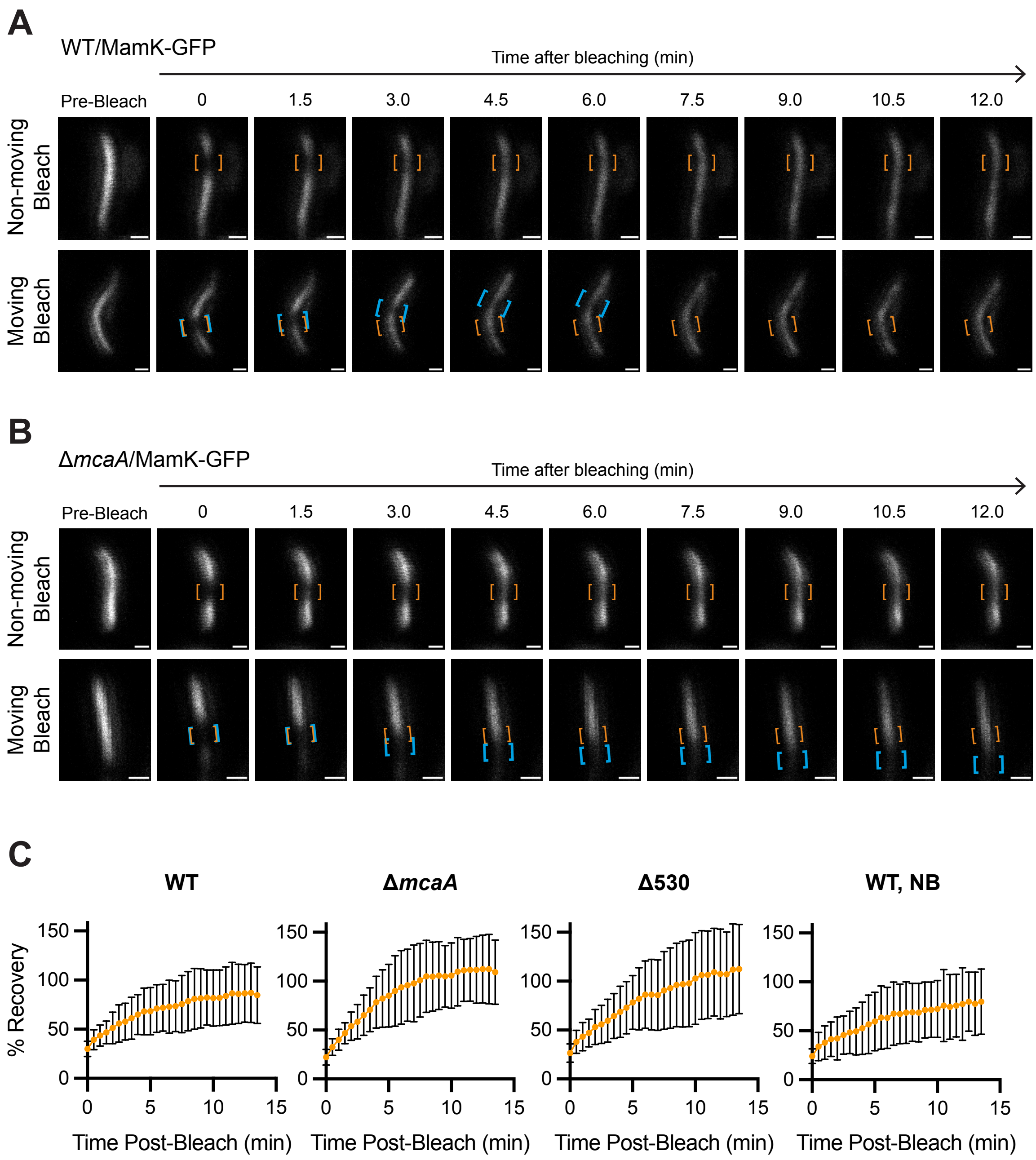


**FIG S11**: MamK-GFP dynamics in WT, Δ*mcaA*, and Δ530. Two representative cells from FRAP of MamK-GFP in WT (**A**) and Δ*mcaA* (**B**). Orange brackets indicate the original bleaching location. Blue brackets track bleach spot movements. The top cell has a bleach spot that does not move while the bottom cell has a moving bleach spot. Scale bar = 0.5μm. (**C**) Percent recovery of MamK-GFP photobleached region in WT, Δ*mcaA*, Δ530, and WT grown in non-biomineralization conditions (WT, NB). The percent recovery is normalized to the whole filament fluorescence at every time point. The percent recovery averages ± standard deviations across all cells are shown. Detailed data can be found in the FRAP source data file.

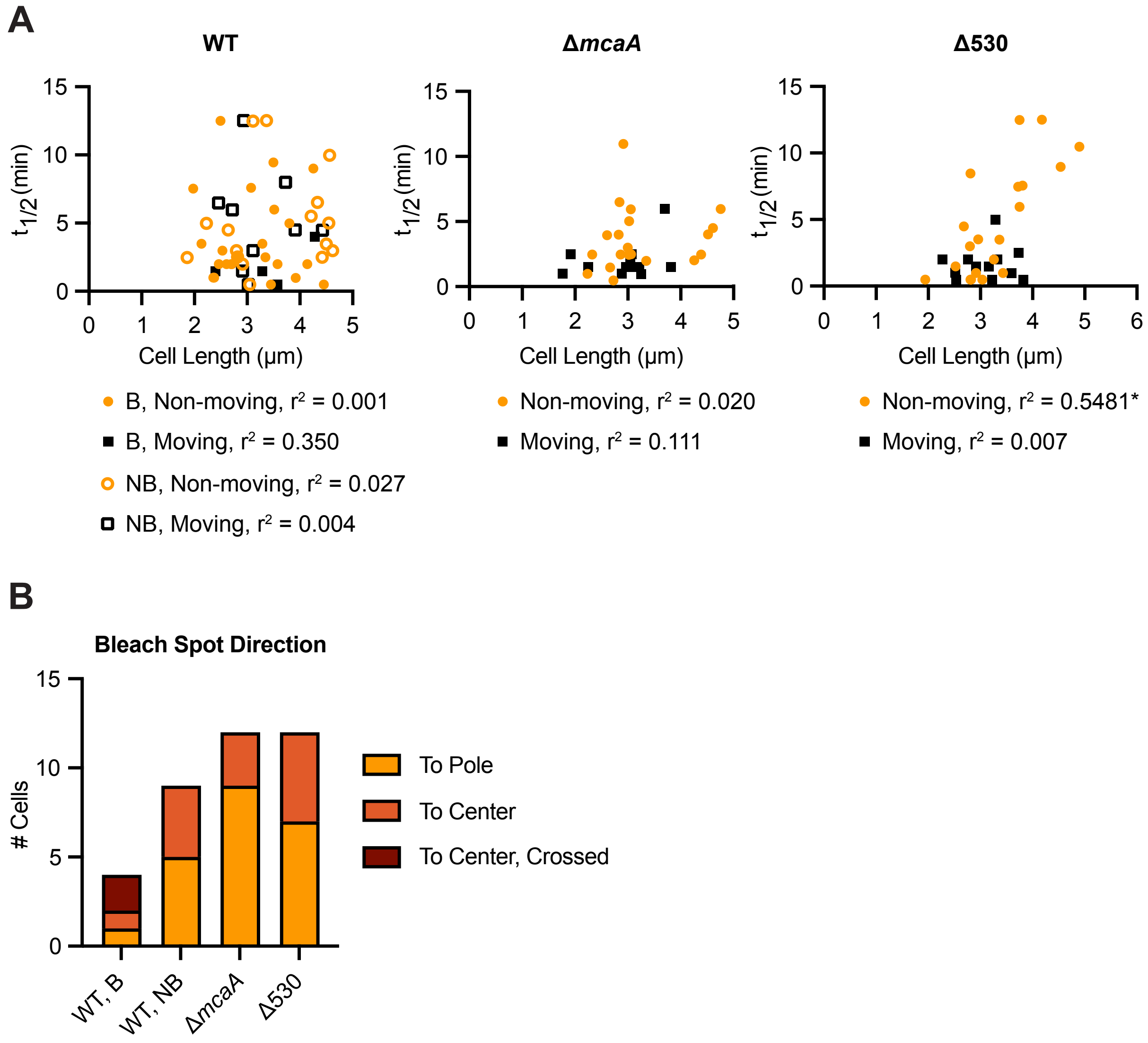


**FIG S12**: MamK-GFP recovery half-times are mostly independent from cell length and bleach spot recovery direction is inconsistent. (**A**) The recovery half-times ($t_{1/2}$) from MamK-GFP FRAP experiments are plotted against the cell length. Each point represents one cell. Yellow circles and black squares represent cells with non-moving bleach spots and with a moving bleach spot, respectively. For WT, filled circles and squares represent cells grown in biomineralization conditions (B) while empty ones represent those grown in non-biomineralization conditions (NB). Cells that did not recover were excluded from the plot. The $r^2$ values and significance generated from simple linear regression are shown (* $P < 0.05$). (**B**) For each cell with a moving bleach spot during the FRAP experiments, the movement direction was categorized as going towards a cell pole or towards the cell center. Cells with bleach spots that moved towards the cell center and crossed through it during the FRAP observation time frames were categorized separately ("To Center, Crossed"). The numbers of cells with each moving direction across each strain and biomineralization condition are shown.

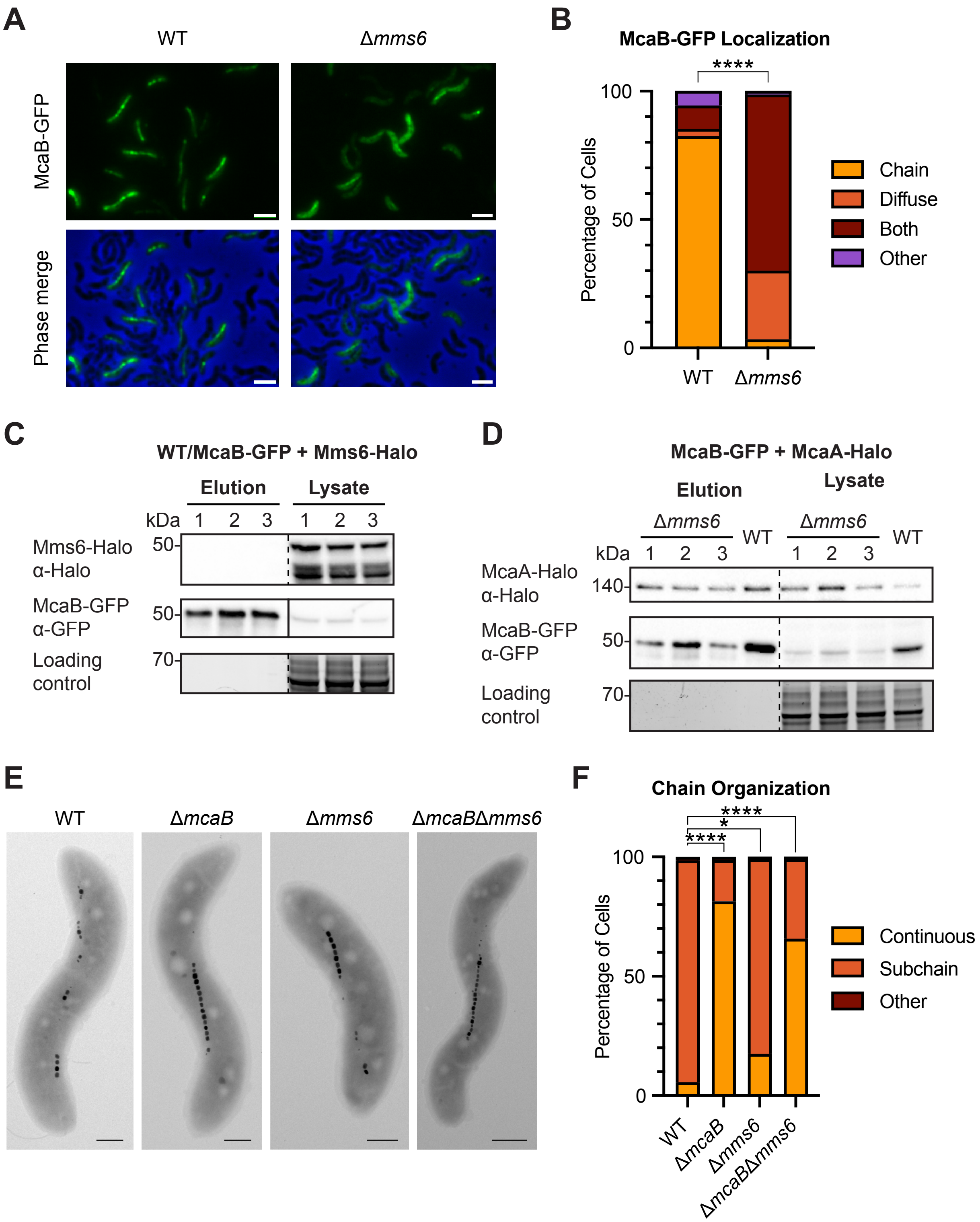
A
WT
Δmms6
McaB-GFP
Phase merge
B
McaB-GFP Localization
****
100
50
0
Percentage of Cells
WT
Δmms6
Chain
Diffuse
Both
Other
C
WT/McaB-GFP + Mms6-Halo
Elution
Lysate
kDa
1 2 3
1 2 3
Mms6-Halo
α-Halo
50-
McaB-GFP
α-GFP
50-
Loading
control
70-
D
McaB-GFP + McaA-Halo
Elution
Lysate
Δmms6
WT
Δmms6
WT
kDa
1 2 3
1 2 3
McaA-Halo
α-Halo
140-
McaB-GFP
α-GFP
50-
Loading
control
70-
E
WT
ΔmcaB
Δmms6
ΔmcaBΔmms6
F
Chain Organization
****
*
****
100
50
0
Percentage of Cells
WT
ΔmcaB
Δmms6
ΔmcaBΔmms6
Continuous
Subchain
Other

**FIG S13**: Mms6 is not required for crystal subchain formation or McaA-McaB interactions. (**A**) Representative cells showing McaB-GFP localization in WT and Δ*mms6* strains. Scale bar = 2μm. (**B**) Quantification of McaB-GFP localization using fluorescent microscopy images. Cells were categorized based on their localization pattern and the y-axis represents the percentage of cells that displayed the indicated localization. WT n = 2316, Δ*mms6* n = 3663. Fisher's exact test was used to determine p-values (**** $P < 0.0001$). (**C**) Representative TEM images of WT, Δ *mcaB*, Δ*mms6*, and Δ*mcaB*Δ*mms6* to show the magnetosome chain organization. Strains were grown anaerobically since AMB-1 strains missing *mms6* makes poor crystals in microaerobic conditions. Scale bar = 0.5μm. (**D**) Quantification of magnetosome chain organization using TEM images. Cells were categorized based on their magnetosome chain phenotype and the y-axis represents the percentage of cells that displayed the indicated chain organization. WT n = 71, Δ*mcaB* n = 146, Δ*mms6* n = 103, Δ*mcaB*Δ*mms6* n = 108. Fisher's exact test was used to determine p-values (* $P < 0.05$, **** $P < 0.0001$). (**E**) Western blots of co-IP elutions and lysates illustrating that Mms6-Halo and McaB-GFP do not interact. (**F**) Western blots of co-IP elutions and lysates illustrating that McaA-Halo interacts with McaB-GFP in Δ*mms6*. WT samples serve as a positive control. In both (E) and (F), GFP-Trap agarose beads were used for the co-IP. Three independent biological replicates are shown and a SDS-PAGE gel serves as a loading control.

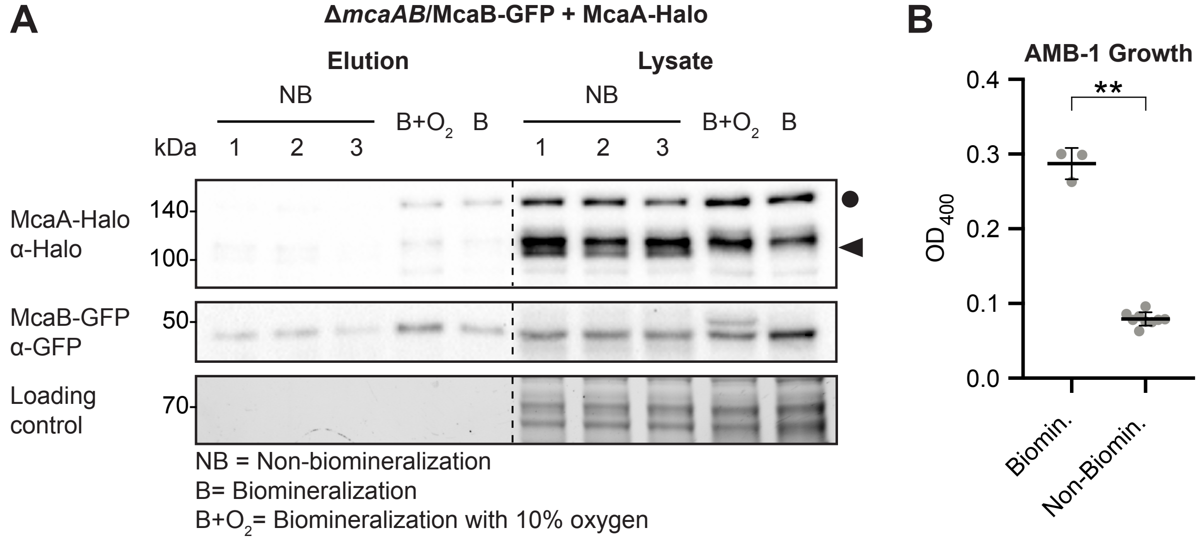


**FIG S14**: McaA and McaB do not interact in non-biomineralizing conditions. (**A**) Western blots of elutions from co-IP with GFP-Trap agarose beads and lysates illustrating that full-length McaA-Halo does not interact with McaB-GFP in the Δ*mcaAB* strain grown under non-biomineralizing conditions (NB). Biomineralization conditions with 10% $O_2$ (B + $O_2$) and normal biomineralization conditions (B) serve as positive controls. The circle and triangle indicate the positions of full-length McaA-Halo and shorter McaA bands, respectively. Three independent biological replicates are shown and the loading control is a stain-free SDS-PAGE gel. (**B**) $OD_{400}$ of 10mL liquid cultures of AMB-1 Δ *mcaAB* expressing McaA-Halo and McaB-GFP microaerobically grown for 2 days in either biomineralization or non-biomineralization conditions. Each point represents an independent culture and the average and standard deviation are shown. Mann-Whitney *U* test was used to determine p-values (** $P < 0.01$).
